# Antagonistic small proteins enable magnesium-dependent tuning of two-component signaling

**DOI:** 10.64898/2026.05.06.723403

**Authors:** Lisa Werland, Gabriele Malengo, Wenting Liu, Julia Rinn, Karim H Kholeif, Timo Glatter, Remy Colin, Victor Sourjik, Jing Yuan

## Abstract

Small proteins are emerging regulators of bacterial virulence via signaling pathways. How multiple small proteins coordinate to control a single pathway remains unclear. Here we show that two antagonistic small proteins, SafA and MgrB, form a magnesium-dependent control module and enhance the signaling sensitivity of the PhoQ/PhoP two-component system in *Escherichia coli*. Leveraging conditions enabling sequential expression of SafA and MgrB, we characterize their regulatory dynamics. Our results reveal magnesium as the central control parameter modulating the abundance and affinities of the small proteins for the sensor kinase PhoQ. This coupling together with the competitive binding of the small proteins enables self-adjustment of the signaling network and enhances its sensitivity to environmental magnesium changes. Functionally, the activity of either small protein influences the ability of enteropathogenic *E. coli* to evade macrophage phagocytosis, linking this regulatory scheme to host interaction. Our findings establish a framework for achieving a high level of input sensitivity through antagonistic regulation in a signaling network.

## Introduction

Small proteins are directly translated from small open reading frames (sORFs) with length cutoffs ranging from 50 to 100 amino acids (*1–5*). Advances in ribosomal profiling, proteomics and bioinformatics tools have uncovered a rapidly expanding collection of small proteins across all three domains of life (*6, 7*). Because of their size, small proteins rarely adopt elaborate folds or exhibit enzymatic activity. Instead, small proteins mostly act through direct physical interactions with larger molecules (*3, 8, 9*). In bacteria, small proteins contribute to various cellular processes including virulence regulation (*3, 4, 10–16*).

The PhoQ/PhoP two-component system (TCS) is considered as the master regulator of the bacterial virulence program in *Salmonella*, *E. coli* and related bacteria (*17, 18*). The system detects host-associated cues and regulates the expression of genes involved in acid resistance, stress response, Mg^2+^ homeostasis, membrane modification and resistance against antimicrobial peptides (AMPs) (Figure 1A) (*19–26*). In enteropathogenic *E. coli* (EPEC), the PhoQ/PhoP system also regulates the expression of the type III secretion system, the effectors, and additional factors contributing to EPEC virulence (*27–31*).

**Figure 1.**
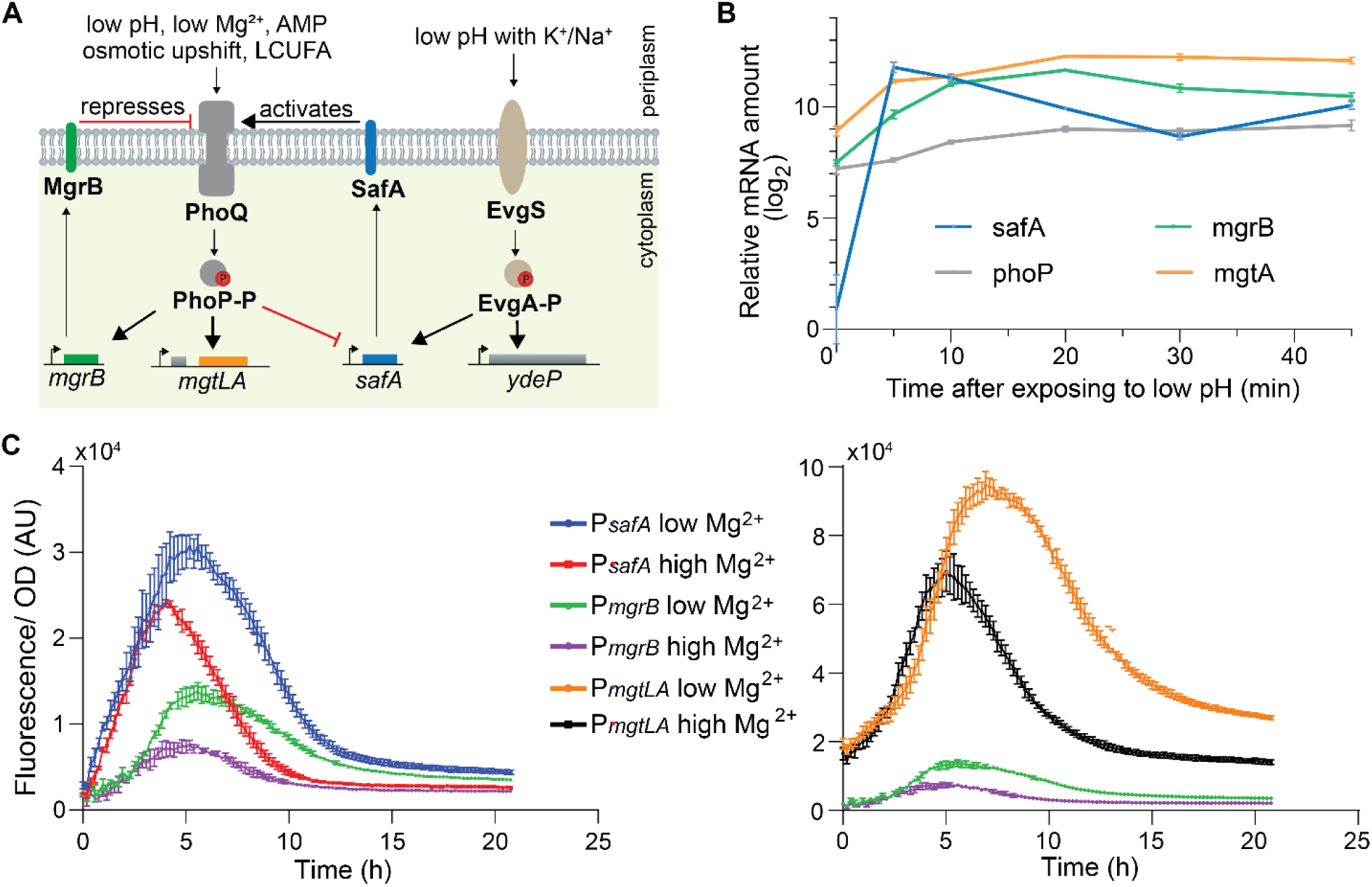
Regulation of PhoQ activity by SafA and MgrB over time. **A**) Schematic overview of the regulation networks of the PhoQ/PhoP and the EvgS/EvgA TCSs. The activation of EvgS by low pH in the presence of alkaline ions leads to the expression of *safA*. The small regulatory protein SafA then activates the PhoQ/PhoP system via the sensor kinase. The phosphorylated PhoP represses *safA* induction acting as a negative feedback, meanwhile it induces *mgrB* expression, which represses PhoQ activity. **B**) The change of the indicated transcripts amounts in *E. coli* MG1655 after exposing to low pH. Briefly, the wild type cells were grown to exponential phase in M9 minimal medium supplemented with potassium at pH 7.0. After switching to pH 5.5, samples were taken at indicated time points followed by total RNA extraction and quantification of *safA*, *mgrB*, *phoP*, and *mgtA* transcripts via RT-qPCR. **C**) The activation profiles of *safA*, *mgrB* and the operon *mgtlA* during the growth of the wild type cells in an EvgS activating medium with low or high magnesium concentrations. GFP fluorescence and optical density (OD) were measured over the course of cell growth in minimal E medium with low (0.1 mM) or high (10 mM) Mg^2+^ at pH 5.5. The ratio of GFP fluorescence to OD_600_ were plotted over time. The P*_mgrB_* dataset is plotted in both charts as the reference. In B and C, data points and error bars represent the mean and standard deviation of three biological replicates. One set of representative results out of three independent experiments is shown.

The PhoQ sensor kinase is a membrane-localized constitutive dimer (*32, 33*). The small membrane protein MgrB interacts with the PhoQ transmembrane (TM) and periplasmic domains, inhibiting PhoQ activity and facilitating a negative feedback loop (Figure 1A) (*34–36*). MgrB prevents PhoQ hyperactivation, mediates partial adaptation and serves as an entry point for redox potential and AMPs (*13, 34, 35*). One main interaction site of PhoQ and MgrB is located in the PhoQ periplasmic domain, which has been proposed to overlap with the putative SafA binding site (*13, 26, 35, 36*). The small membrane protein SafA activates PhoQ kinase activity by accelerating the autophosphorylation (*37, 38*). The C-terminal region of SafA interacts with the PhoQ periplasmic domain, responsible for the activation (*26, 38*). SafA is expressed in an operon with the downstream *ydeO* from a promoter regulated by the EvgS/EvgA TCS under mild acidic conditions. It connects the PhoQ/PhoP and EvgS/EvgA systems and increases *E. coli* survival under acidic conditions at a pH comparable to the intragastric environment (*37, 39*). Additionally, PhoP lowers the expression of *safA* as shown by the elevated expression from the *safA* promoter in a Δ*phoP* strain (*40*) (Figure 1A).

TCSs are known to form intricate signaling network. Although the regulation of PhoQ activity by MgrB and SafA has been characterized individually, how the antagonistic regulations are coordinated and the impact on bacterial adaptation remain unclear. This is partly due to the overlapping abilities of EvgS and PhoQ in sensing low pH, which obscures the identification of the sensor kinase responsible for initiating specific signaling cascades and complicates the analysis of gene regulation. In this study, we identify low pH conditions that directly activate EvgS but activate PhoQ indirectly via SafA. Under these conditions, we characterize the expression dynamics of MgrB and SafA and their interaction with PhoQ *in vivo* via GFP reporters, Förster resonance energy transfer (FRET) approaches, and mathematical modeling. The small proteins exhibit divergent expression profiles under low pH conditions. Higher magnesium enhances the affinity of MgrB to PhoQ and promotes PhoQ inhibition, whereas lower magnesium leads to an increase of *safA* expression, a preferred binding of PhoQ to SafA, promoting PhoQ activation. Our mathematical model recapitulates the regulatory dynamics, showing that this magnesium-dependent competitive binding of the small proteins enhances the sensitivity of the signaling network towards magnesium changes. Furthermore, we demonstrate that the regulation of MgrB and SafA is physiologically important. Deletion of *mgrB* or increasing the expression of *safA* significantly enhances EPEC evasion of macrophage phagocytosis.

## Results

### Regulation of PhoQ activity by SafA and MgrB over time

PhoQ is activated by mild acidic pH in *E. coli* (*35*). However, it was unclear whether this activation is direct or through the small protein SafA, since low pH could also activate the EvgS sensor kinase (*41*) (Figure 1A). To clarify this, we constructed a Δ*safA* strain in *E. coli* MG1655, and monitored PhoQ activity using a GFP reporter (P*_mgtLA_*-*gfp*) under mild acidic conditions. The EvgS activity was followed using a different GFP reporter (P*_ydeP_*-*gfp*) (*42*). At pH 5.5, we observed an increase of GFP fluorescence for EvgS activity, when cells were grown in potassium-supplemented M9 medium, confirming its activation (Figure S1A). This response required potassium consistent with previous observations (*41*). PhoQ activity increased at low pH in the wild type but not in the Δ*safA* strain, indicating SafA is required for PhoQ activation (Figure S1B) under this growth condition.

We also followed EvgS and PhoQ activities in other growth media (minimal A and minimal E) with a different set of reporters (P*_mgrB_*-*gfp* and P*_safA_*-*gfp* for PhoQ and EvgS activities, respectively). Reproducibly, we observed that fluorescence increase of the PhoQ activity reporter was only detected in the wild type but not in the Δ*safA* strain at pH 5.5 (Figure S2). These results strongly suggest that under minimal medium growth conditions, low pH activates PhoQ via EvgS and SafA in *E. coli*.

Leveraging on this sequential activation of the EvgS/EvgA and PhoQ/PhoP systems under low pH conditions, we studied how SafA and MgrB modulate signal propagation and PhoQ activation. To do so, we monitored the transcript abundance for *safA* and genes in the PhoP regulon over time. The transcript amounts were quantified with RT-qPCR at different time points after exposing the cells to low pH during exponential growth (Figure 1B). We detected a pronounced increase of *safA* transcripts upon low pH stimulation, with mRNA levels rising from basal background to a maximum within five minutes. The *safA* transcript level was tapered off slightly and stabilized around 40 minutes at a level comparable to the *mgrB* transcript (Figure 1B and S3). Regulated by the PhoQ/PhoP system, *mgrB*, *phoP,* and *mgtA* had markedly elevated basal transcript amounts at time 0, with their levels reaching the maximum much later (in 20 min), and stabilizing around the same time as *safA* (around 40 min). This relatively slow activation of PhoP regulated genes is likely due to the time needed for SafA to be expressed, inserted into the inner membrane, and activate PhoQ. Notably, the increase of the *phoP* transcript at 5 min post-activation coincided with a decline in *safA* transcript levels, consistent with previous observations that PhoP represses *safA* expression (*40*).

To assess the influence of PhoQ activation dynamics by MgrB and SafA for a longer period across different growth phases, we used two GFP reporters (P*_safA_*-*gfp* and P*_mgrB_*-*gfp*) to follow the activation of EvgS and PhoQ respectively. The two promoter reporters also served as indicators for SafA and MgrB production during the analysis. As a read-out, we measured GFP fluorescence over time in MG1655 grown at a mild acidic condition (pH 5.5) in the presence of low (0.1 mM) and high (10 mM) magnesium concentrations (Figure 1C, left).

In line with the transcript analysis, the fluorescence from the P*_safA_*-*gfp* reporter increased earlier than that from P*_mgrB_*-*gfp* in the wild type strain, confirming the sequential activation of the EvgS/EvgA and PhoQ/PhoP TCSs under this growth condition. The *safA* promoter fluorescence also peaked slightly earlier and at a higher value than that of the *mgrB* promoter reporter. Magnesium had a marked influence on both reporter activities. Lowering the Mg²⁺ concentration to 0.1 mM Mg²⁺ resulted in an enhanced activation for P*_safA_*, with its maximum observed during early exponential growth phase at a fluorescence/OD value around 30,000 (Figure 1C). Reduced magnesium acted as an activating condition for PhoQ, increasing P*_mgrB_* activation as expected (*43*), but with a weaker overall signal strength when compared with P*_safA_*.

The activation profiles of *safA* and *mgrB* were also distinct, which was easier to comprehend when plotting the normalized fluorescence (fluorescence/OD) against the growth (Figure S4). The P*_safA_* reporter fluorescence accumulated during the lag phase, which dropped as cells entered the exponential phase, suggesting a fast but possibly temporal activation. This activation appeared weaker under high magnesium conditions, as indicated by a lower maximum and a sharper decay of the fluorescence signal during exponential growth. The activation of P*_mgrB_* began later, with GFP fluorescence increasing in the lag phase and relatively more persistent during exponential growth, suggesting that genes in the PhoP regulon were activated through SafA as well as through low magnesium in these growth phases. In addition, we monitored the GFP fluorescence of a different promoter reporter (P*_mgtLA_*-*gfp*) regulated by PhoP (Figure 1C, right and S4). As expected, this promoter showed higher strength but comparable dynamics as P*_mgrB_*. To assess the possible effect of magnesium alone on reporter expression, we monitored constitutive *mneongreen* expression under high (10 mM) and low (0.1 mM) Mg²⁺ conditions. Fluorescence remained comparable throughout growth, indicating minimal effects of Mg²⁺ alone on the reporter signal (Figure S4D).

Previous results suggest that low magnesium activates PhoQ and induces *phoPQ* expression (*21*), which leads to an enhanced repression of P*_safA_* (*40, 43*). Surprisingly, we consistently observed higher P*_safA_* induction under low magnesium conditions, indicating the presence of additional regulatory factors controlling *safA* expression. As this regulation appeared to be magnesium-dependent, we sought to investigate the potential impact of magnesium on proteins within the EvgS/EvgA and PhoQ/PhoP signaling network.

### Magnesium enhances MgrB binding to PhoQ

High magnesium promotes PhoQ to adopt an inactive conformation, which might affect its affinity to the small regulatory proteins. To determine whether magnesium concentration affects PhoQ/MgrB and PhoQ/SafA interactions, we used established FRET approaches, which exploit energy transfer between donor and acceptor fluorescent proteins at distances smaller than 10 nm (*36, 44*), to monitor these interactions *in vivo* across a range of magnesium concentrations. Plasmids expressing PhoQ-mNeonGreen, mCherry-MgrB, and mCherry-SafA were used to produce the two FRET pairs (PhoQ-mNeonGreen with mCherry-MgrB, and PhoQ-mNeonGreen with mCherry-SafA) in *E. coli*. To prevent any interference from the untagged genomic copy, a triple deletion strain, Δ*phoPQ*Δ*mgrB*Δ*safA*, was constructed and transformed with the FRET plasmids. Cells expressing the FRET pairs were analyzed using stimulus-dependent ratiometric FRET. We excited mNeonGreen and recorded the emissions over time in both the mNeonGreen and mCherry channels. A higher ratio of mCherry vs. mNeonGreen emissions (the red/green ratio) indicates an increase in energy transfer, therefore a stronger interaction between the tagged proteins, and *vice versa*.

When monitoring the binding of MgrB to PhoQ with various magnesium concentrations, we observed an increase in the red/green ratio with higher Mg^2+^ concentrations in the buffer (Figure S5). Inversely, the ratio decreased to the basal level when the Mg^2+^ concentration was reduced from 10, 1, and 0.1 mM back to 0 mM. A similar effect was detected with other divalent cations including Ca^2+^, Mn^2+^, and Zn^2+^ at different concentration ranges (Figure S5). We then tested these cations at lower and more physiologically relevant concentrations (Figure 2A). Variations of Mg^2+^, Ca^2+^, Mn^2+^, and Zn^2+^ reproducibly resulted in changes in the red/green ratio but not with Fe^2+^ (Figure 2A). In contrast, no ratio changes were detected for SafA/PhoQ interactions under these conditions (Figure 2B), suggesting that magnesium, calcium, manganese, and zinc divalent cations enhance PhoQ interaction with MgrB but not SafA at the tested concentrations.

**Figure 2.**
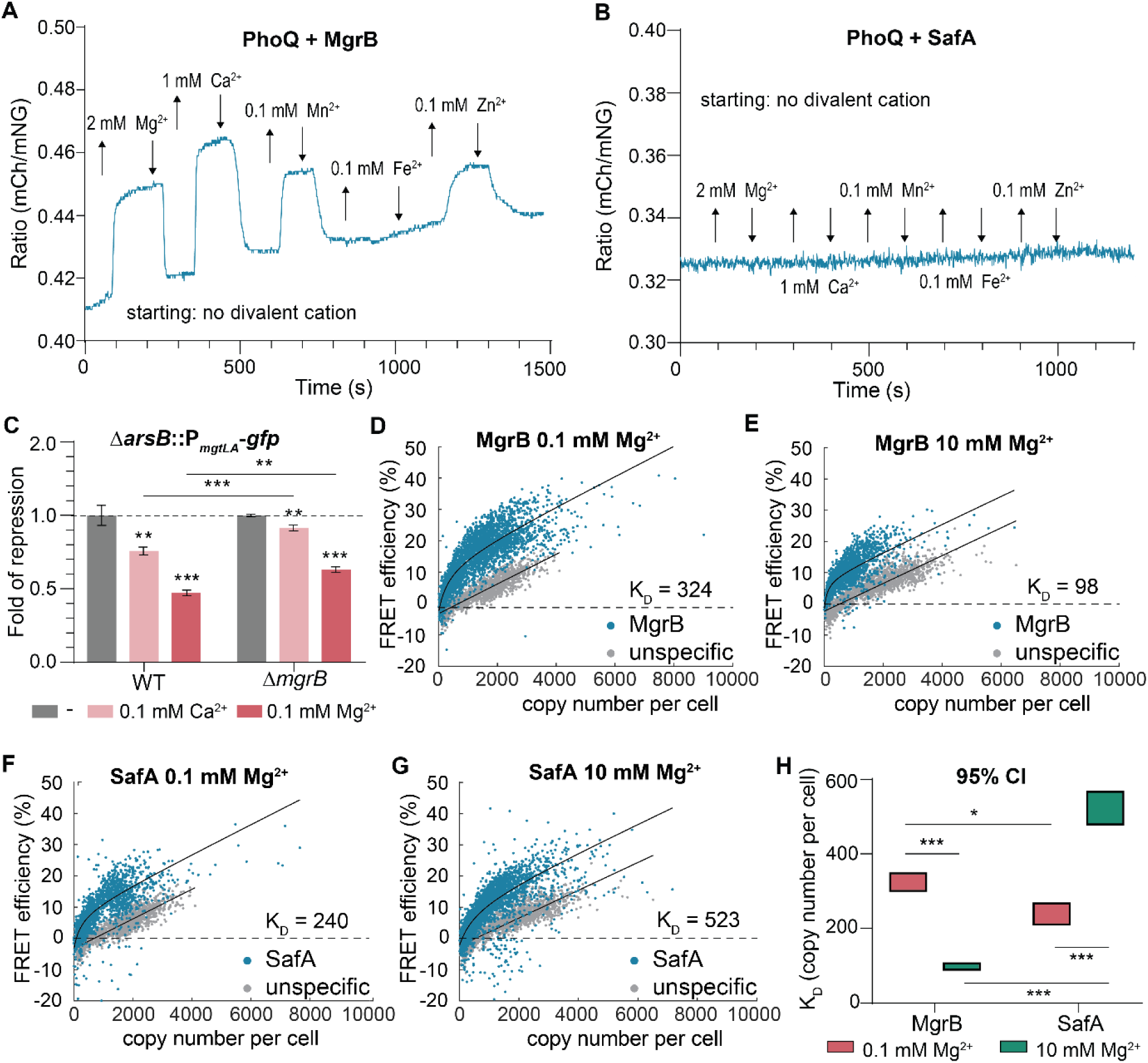
Magnesium modulates the interaction of PhoQ to MgrB and SafA. Ratiometric FRET of PhoQ-mNeonGreen with (**A)** mCherry-MgrB and (**B**) mCherry-SafA. The FRET pairs were expressed in MG1655 Δ*phoPQ*Δ*mgrB*Δ*safA* and the cells were immobilized inside a flow chamber that was kept under a constant flow of buffer. The fluorescence intensity of mNeonGreen and mCherry was acquired by fluorescence microscopy. The ratio of mCherry intensity over mNeonGreen intensity was plotted over time. Divalent cations were added (upward arrow) and removed (downward arrow) at indicated concentrations and time points by exchanging the buffer. **C)** Repression of PhoQ by divalent cations in presence (WT) and absence of MgrB (Δ*mgrB*). PhoQ activity was monitored using a GFP reporter integrated into the genome (Δ*arsB*::P_mgtLA_-*gfp*). Strains were grown in modified minimal A medium supplemented with the indicated concentrations of divalent cations. Error bars represent the mean and standard deviation of the median fluorescence intensity of 30,000 cells of three biological replicates per condition. One representative result out of three independent experiments is shown in A, B and C. **D-G)** Binding affinity of MgrB and SafA to PhoQ at low and high [Mg^2+^]. FRET efficiency was determined for single cells expressing PhoQ-mNeonGreen and mCherry-MgrB (blue in D and E) or mCherry-SafA (blue in F and G). The FRET measurement between PhoQ-mNeonGreen and MalFTM1 (grey) was used as a control for non-specific binding. FRET efficiency of each cell is plotted as a function of the mCherry copy number per cell as single points. The K_D_ was estimated by fitting the data according to Michaelis-Menten kinetics accounting for the non-specific binding deduced from MalFTM1. **H**) The calculated K_D_ values in the 95% confidence interval (CI). P values were calculated using a Student’s t-test (ns: P>0.05; *: P < 0.05; **: P < 0.01; ***: P < 0.001).

To assess whether the observed binding changes of MgrB and PhoQ has functional consequences, we compared PhoQ activity in strains with and without MgrB under growth conditions lacking or supplemented with either Ca²⁺ or Mg²⁺ using the GFP reporter for PhoQ (P*_mgtLA_*-*gfp*). PhoQ activity in general was higher in the absence of MgrB in both growth conditions due to lack of the repressor MgrB (Figure S6). After normalization, MgrB in presence of Mg^2+^ and Ca^2+^ resulted in a stronger inhibition of the PhoQ activity (Figure 2C), suggesting that the enhancement in PhoQ/MgrB interaction by magnesium and calcium ions has a functional effect and leads to an increased inhibition of PhoQ. For Mn^2+^ and Zn^2+^, we did not detect any impacts on MgrB inhibition of PhoQ nor inhibition of PhoQ by the cation themselves (Figure S6), consistent with a previous report (*45*). Taken together, our data support the hypothesis that Mg^2+^ and Ca^2+^ stabilize PhoQ in a conformational state, which allows better binding and further repression of PhoQ by MgrB.

### The binding affinities of SafA and MgrB to PhoQ

To compare and quantify the bindings of the small proteins to PhoQ, we used single-cell acceptor photobleaching FRET (*46, 47*) to estimate the dissociation constants (K_D_s). FRET efficiency was quantified at the single-cell level under controlled induction conditions, in which the concentration of PhoQ–mNeonGreen was maintained constant, while the levels of mCherry–MgrB or mCherry–SafA were systematically titrated across a broad concentration range. As a quantification of protein abundance, we converted fluorescence of mCherry to intracellular copy number by using purified mCherry with known concentrations as reference (Figure S7). To estimate the contribution to FRET from non-specific protein-protein interactions, we fused mCherry to the first TM helix of MalF (MalFTM1), which was indicated as a non-binder of PhoQ in a previous study (*36*). MalFTM1 showed a linear increase in FRET efficiency with increasing copy numbers per cell, demonstrating the increase in energy transfer and its non-specific interactions with PhoQ (Figure 2D-G gray). The negative FRET values at the very low concentration range were likely due to a low signal-to-noise level and possible photo-damaging of the donor fluorophore.

Unlike the non-specific binding control, MgrB and SafA showed hyperbolic increase in energy transfer to PhoQ with increasing copy numbers of the small proteins per cell (Figure 2D-G blue). Larger deviations in FRET measurements were observed at higher amounts of the mCherry-tagged small proteins, likely due to a higher variation in protein stability and membrane insertion. The saturated energy transfer from MgrB to PhoQ slightly varied under different magnesium conditions, possibly due to the PhoQ or MgrB conformational change induced by magnesium. We fitted the single-cell FRET measurements with Michaelis-Menten kinetics accounting for the non-specific binding deduced from MalFTM1, and calculated the K_D_ values of MgrB and SafA to PhoQ. At the low magnesium condition (0.1 mM), SafA showed slightly higher affinity to PhoQ than MgrB with a smaller K_D_ value (240 vs. 324 copies per cell, Figure 2D and F). Higher magnesium showed a divergent effect on the binding affinities of the two small proteins. The calculated K_D_ of MgrB reduced about 3-fold (from 324 to 98 copies per cell), indicating an enhancement in PhoQ binding, consistent with our previous observations at the population level (Figure 2A). In contrast, the calculated K_D_ of SafA increased about 2-fold (from 240 to 523 copies per cell) under the high magnesium condition, suggesting a reduction in PhoQ binding (Figure 2F and G). The single-cell FRET measurements of PhoQ and SafA generally showed more variation than that of PhoQ and MgrB (possibly due to more variations in expression) resulting in larger deviations in the 95 % confidence values (Figure 2H).

Collectively, these results demonstrate that SafA binds to PhoQ slightly better than MgrB at low magnesium concentrations with the K_D_ ^SafA^ in the range of 200 to 300 copies per cell. The binding preference of PhoQ shifts to MgrB in high magnesium, with the K_D_ ^MgrB^ reduced to about 100 copies per cell. Our single-cell FRET data quantitatively supports the notion that magnesium adds another layer of regulation on the PhoQ activity through modulating the binding affinities of the small regulatory proteins. Lower magnesium leads to high PhoQ activity by favoring the interaction of PhoQ to its activator SafA and by direct activation. Higher magnesium results in low PhoQ activity by favoring the interaction of PhoQ to its inhibitor MgrB and direct repression.

### MgrB and SafA compete for interaction with PhoQ

Previous reports suggest that MgrB and SafA may partially share binding sites on PhoQ (*26, 35*). To test whether MgrB and SafA compete for the interaction with PhoQ, we measured the energy transfer of PhoQ-mNeonGreen to mCherry-MgrB in the presence of SafA and *vice versa*. The function of mCherry-tagged small proteins showed no significant change or only a slight decrease compared to the untagged versions, indicating a minimal impact of the N-terminal mCherry tag (Figure S8). Additionally, FLAG tags were added to facilitate protein quantification.

Using the construct of P_IPTG_-*mcherry-mgrB*-P_con_-*safA* (Figure 3A), IPTG-induced levels of mCherry-MgrB were lower than the constitutive SafA levels (Figure 3B, left). We then measured the energy transfer from PhoQ to MgrB without or with SafA. The additional expression of SafA did not markedly affect the ratio of mCherry to mNeonGreen fluorescence (right Y-axis in Figure 3B), indicating the acceptor to donor ratios were comparable in both setups. The FRET results showed significant reduction in energy transfer when SafA was present, suggesting that SafA competes with MgrB for PhoQ binding at this condition.

**Figure 3.**
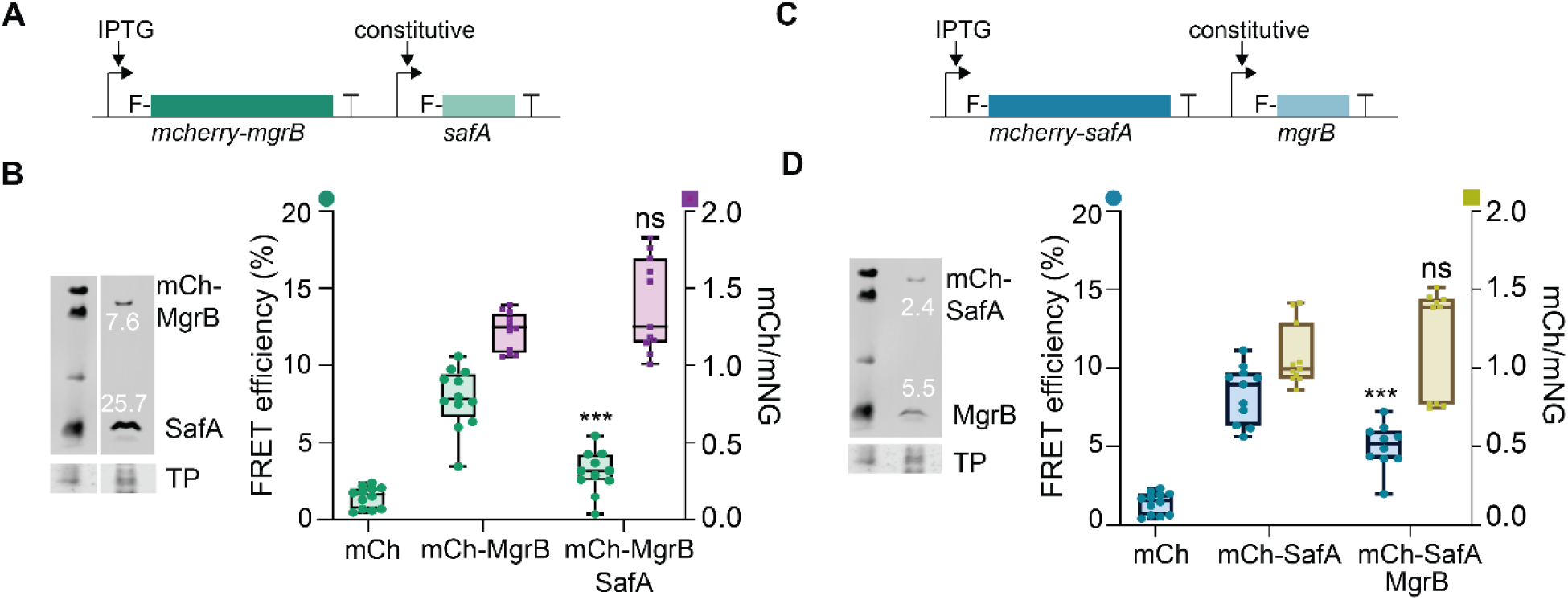
Competition of MgrB and SafA for binding to PhoQ. Plasmid constructs are shown for assessing the influence of SafA on PhoQ-MgrB interaction (**A**) and that of MgrB on PhoQ-SafA interaction (**C**). Expression of a mCherry-fusion was induced with 10 µM IPTG. The expression of the FLAG-tagged small protein was constitutive. F stands for FLAG-tag. **B**) *E. coli* MG1655 Δ*phoPQ* Δ*mgrB* Δ*safA* expressing PhoQ-mNeonGreen and mCherry MgrB (mCh-MgrB) without or with SafA expressed from the plasmid described in A, were grown for acceptor photobleaching FRET. The left Y-axis represents FRET efficiency (green data points). The right Y-axis is for the ratio of mCherry over mNeonGreen (purple data points), which serves as an indicator of acceptor/donor expression. Expression of mCh-MgrB and SafA was quantified by western blot analysis. The numbers below/above the bands indicate the intensity of the band normalized to the total protein (arbitrary units). **D**) Acceptor-photobleaching FRET of PhoQ-mNeonGreen and mCherry-SafA (mCh-SafA) without or with MgrB expressed from the plasmid described in C. The left Y-axis represents FRET efficiency (blue data points). The right Y-axis is for the ratio of mCherry over mNeonGreen (chartreuse data points), which serves as an indicator of acceptor/donor expression. The FRET efficiency of three biological replicates with each four-measurement points per sample is plotted as box-plots in B and D. One representative western blot out of three biological replicates is shown in B and D. P values were calculated using a Student’s t-test by comparing the values in B (mCherry-MgrB with to without SafA competition) and D (mCherry-SafA with to without MgrB competition). ns: P>0.05; *: P < 0.05; **: P < 0.01; ***: P < 0.001.

In parallel, we tested whether MgrB interfered with SafA binding to PhoQ using the same approach (Figure 3C and D). Using the construct of P_IPTG_-*mcherry-safA*-P_con_-*mgrB*, IPTG-induced levels of mCherry-SafA were roughly two times less than the constitutive MgrB levels (Figure 3D, left). Under this condition, the FRET efficiency from PhoQ-mNeonGreen to mCherry-SafA decreased significantly when compared to that in the absence of MgrB, indicating that MgrB can compete with SafA as well. Taken together, our FRET data support the hypothesis that MgrB and SafA compete for binding to PhoQ.

### Mathematical model of the EvgS/EvgA and PhoQ/PhoP signaling network

We modeled the EvgS/EvgA and PhoQ/PhoP signaling network mathematically to simulate its dynamics in the presence of low and high magnesium concentrations. Low pH as the initial stimulation activated EvgS/EvgA, which then activated the PhoQ/PhoP system through SafA. The model combined our findings with previous knowledge by incorporating the dissociation constants of MgrB and SafA estimated above (Figure 2D-G) as well as their competitive binding to PhoQ (Figure 3). The model also assumes that: i) the small proteins do not affect the binding of Mg²⁺ to PhoQ; ii) the binding of small protein to PhoQ dimer follows 1:1 ratio; iii) PhoP-P binds to *safA* promoter non-competitively with EvgA-P; iv) PhoQ concentration is always proportional to PhoP; and vi) proteins are diluted by growth. We modeled the dynamics of SafA and MgrB production during a lag phase followed by an exponential growth phase, each lasting five doubling times (t_1/2_).

Under a low magnesium condition (blue solid lines in Figure 4A and S9), both small proteins showed a sharp increase in the lag phase followed by a decay in the exponential growth phase, with the simulated SafA concentration changing at a faster rate than MgrB, consistent with our experimental data (Figure 1C and D). Under a high magnesium condition (red dash lines in Figure 4A and S9), the simulated MgrB concentration was lower due to the high magnesium repression of PhoQ. However, the simulated SafA concentration increased due to the reduced PhoP-P repression, contradicting the experimental results, even after incorporating the magnesium-dependent binding of the small proteins to PhoQ.

**Figure 4.**
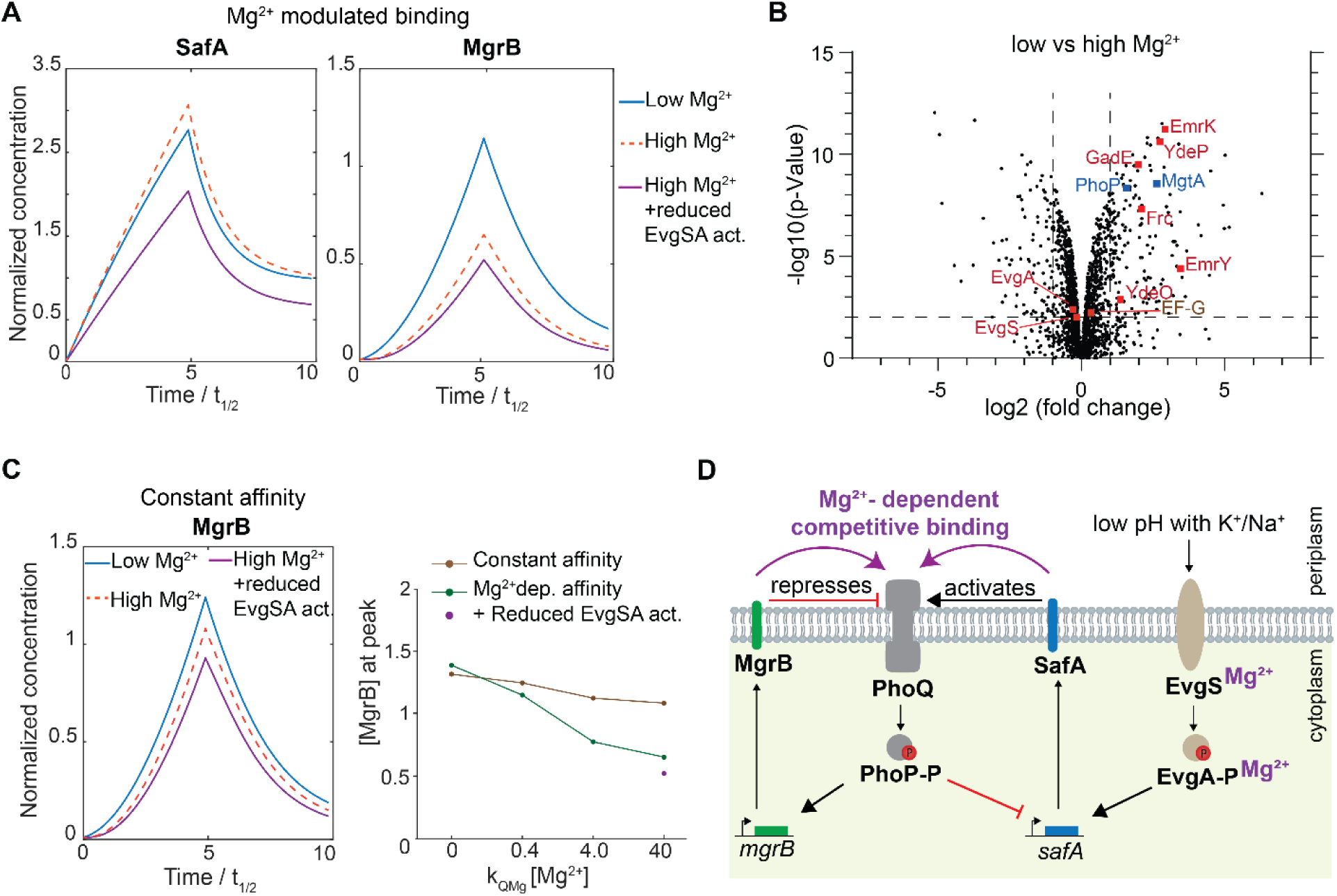
EvgS and EvgA serve as additional entry points for magnesium into the signaling network. **A**) Mathematical simulations of the dynamics of SafA and MgrB concentrations in high and low magnesium conditions upon stimulation with low pH. **B**) Comparison of the *E. coli* proteome grown at low and high magnesium concentrations with a low pH stimulation. Specifically, MG1655 cells were grown in minimal E medium with low (0.1 mM) and high (10 mM) magnesium at pH 5.5 to exponential phase. Cells were then harvested, washed and followed by total protein extraction and proteomic analysis. Data represents the mean values of four biological replicates. **C**) Mathematical simulation of the dynamics of MgrB concentrations omitting magnesium-dependent binding of the small proteins to PhoQ (left). In the right chart, the peak concentration of MgrB as an indicator of PhoQ/PhoP activity is plotted over the magnesium concentration times its binding affinity to PhoQ (k_QMg_). **D**) Schematic summary of the regulation networks of the PhoQ/PhoP and the EvgS/EvgA TCSs with the magnesium influenced nodes highlighted in purple.

One way to reconcile the simulation with the experimental results was to introduce a magnesium-dependent modulation factor for EvgS/EvgA activity (purple solid lines in Figure 4A and S9). Specifically, when we assumed lower EvgS/EvgA activity (65%) under high magnesium conditions; the simulated overall SafA concentration became lower than under low magnesium conditions. These results suggest the presence of a magnesium-dependent input that enters the signaling network through EvgS/EvgA.

To test the prediction by the mathematical simulations, we performed proteomic analysis of the wild type cells grown in a low pH medium (pH 5.5) in the presence of low and high magnesium conditions (0.1 and 10 mM, respectively). Cells were harvested during the exponential growth phase, followed by total protein extraction and mass spectrometry analysis. Under low magnesium conditions, we observed a 2-to 11-fold increase in the levels of proteins regulated by EvgA (Figure 4B and S10). These included YdeO, GadE and YdeP, proteins involved in acid resistance, EmrK and EmrY, a drug efflux pump, and Frc, a formyl-CoA transferase. We did not detect MgrB or SafA due to their small size and hydrophobicity, a common problem for small membrane proteins (*5, 9*). Since *ydeO* is in the same operon and downstream of *safA*, the increase of YdeO abundance suggests the increase of SafA at low magnesium, consistent with our GFP reporter results. The levels of EvgS and EvgA on the other hand did not show marked difference, comparable to that of the EF-G control, suggesting that this magnesium dependent change of EvgA regulon expression is likely due to the change in the phosphorylation level of EvgA and the activity of EvgS instead of EvgS/EvgA abundance. Low magnesium activates PhoQ (*21*). As expected, we observed increased levels of proteins regulated by PhoP, such as MgtA, a magnesium transporter (Figure 4B and S10). These results confirm our model’s prediction that alongside with PhoQ, EvgS and EvgA serve as additional entry points for magnesium into the signaling network.

Magnesium-dependent binding of MgrB and SafA to PhoQ added another layer of regulation in this signaling network. To visualize the benefit of this magnesium-dependent affinity, we additionally simulated a modified model where the binding affinities of SafA and MgrB to PhoQ were set constant, all else being unchanged. In this case, although SafA expression was only marginally affected (Figure S9), MgrB expression, as an indicator of PhoQ activity, showed much less variation at different magnesium concentrations (Figure 4C left and S9). Indeed, in our model, when concentrations of SafA (*a*) and MgrB (*b*) are larger than the dissociation constant, the fraction of active PhoQ depends only on the ratio (*a* / K_D_ ^SafA^) / (*b* / K ^MgrB^) (see Methods), which can only vary as a function of the magnesium concentration via the dissociation constants (K_D_ ^SafA^ and K ^MgrB^). Therefore, the magnesium-modulated binding of the small proteins increases the sensitivity of PhoQ to magnesium changes as shown in our mathematical model (Figure 4C, right).

Taken together, our data support that magnesium as the central control element enters the signaling network by reshaping the equilibria of SafA and MgrB regulatory dominance, and by modulating PhoQ and EvgS/EvgA activities (Figure 4D). This multilayered regulatory system enables self-tuning of the signaling network and enhances sensitivity to magnesium changes.

### MgrB and SafA influence EPEC evasion from macrophages

To evaluate whether the regulation of PhoQ by MgrB and SafA has physiological consequences for pathogenic *E. coli*, we investigated the impact of deletions of *mgrB* and *safA* on evasion of an EPEC strain (E611) from macrophages. The wild type and knockout strains (Δ*mgrB* and Δ*safA*) were grown in minimal E medium at pH 5.5 to log phase under low magnesium (0.1 mM), and then incubated with differentiated human THP-1 macrophages at a multiplicity of infection (MOI) of 100. After two-hour incubation, extracellular bacteria were killed with gentamicin. The proportion of internalized bacteria was quantified by determining colony-forming units (CFUs) and normalizing them to the initial bacterial inoculum (Figure 5A). The cytotoxicity of bacterial infection on macrophages was tested in parallel (Figure S11). Compared to the wild type, we detected a significant reduction of engulfment of Δ*mgrB* cells by macrophages. The Δ*safA* strain showed only a marginally increased uptake by macrophages (Figure 5A, S12).

**Figure 5.**
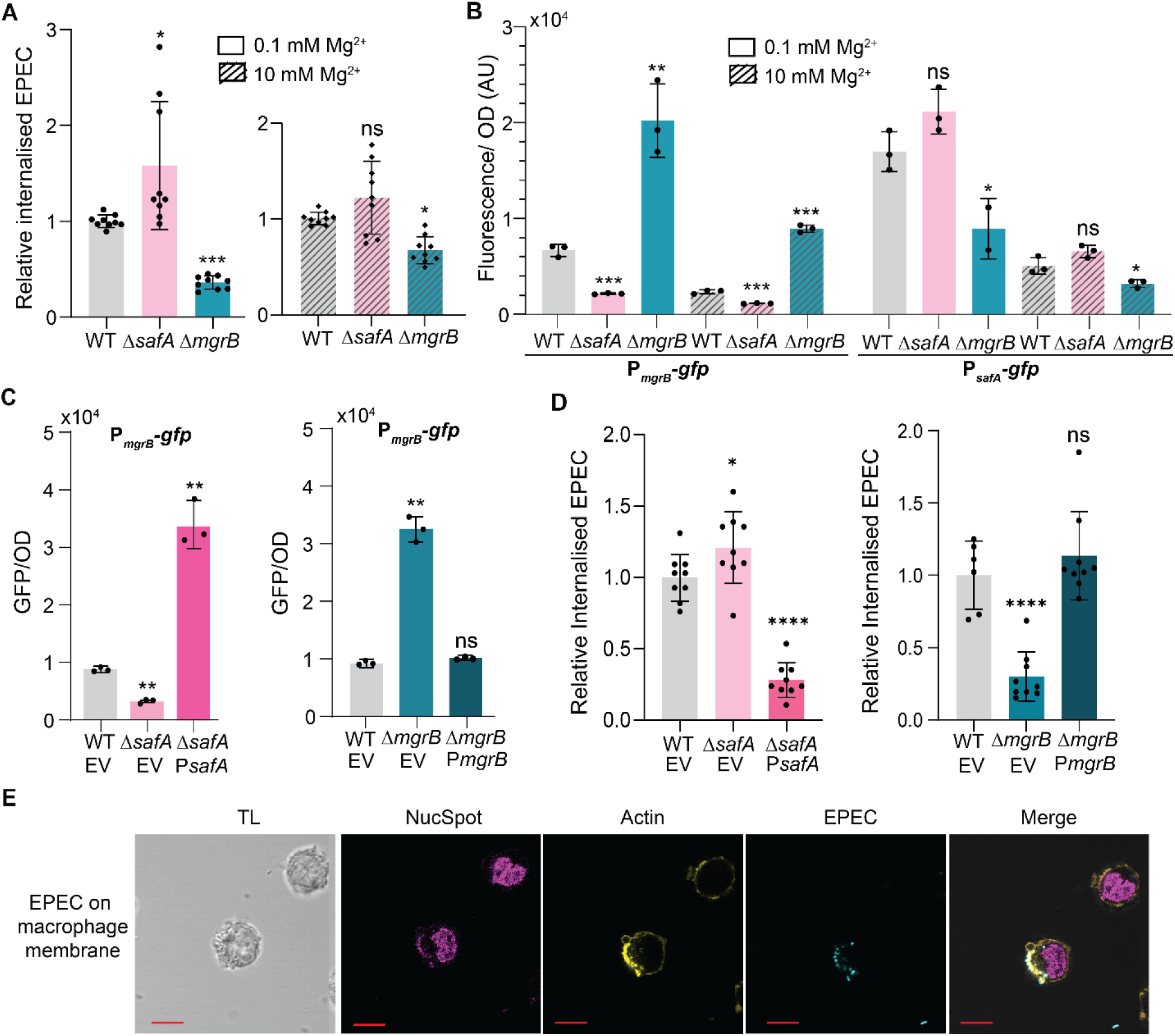
MgrB and SafA influence EPEC evasion from macrophage phagocytosis. **A)** Internalization of EPEC E611 strains in macrophage infection assays. EPEC strains were grown to log phase in minimal E medium at pH 5.5 with indicated Mg²^+^ concentrations, then incubated with macrophages at an MOI of 100 in RPMI medium for 2 hrs. Internalized EPEC were quantified and normalized to the wild-type strain. Data are shown as bar charts (mean ± SD, n = 9 replicates). **B)** Expression of GFP reporters in EPEC strains. EPEC E611 wild type, Δ*safA* and Δ*mgrB* cells carrying the respective reporter plasmids were grown in minimal E medium at pH 5.5 with indicated Mg^2+^ concentrations. GFP fluorescence divided by OD_600_ is plotted for a time point within the exponential growth phase used for infection. **C**) The complementation of Δ*safA* and Δ*mgrB* with the respective plasmids. E611 wild type, Δ*safA* and Δ*mgrB* cells carrying empty vector (EV), pKHK P*_safA_*-*safA* and pBAD33 *mgrB*, respectively. The GFP reporter assay was performed as in B. The growth medium (in C right) was supplemented with 0.15% arabinose to induce *mgrB* expression. **D**) Internalization of the E611 complementated strains by macrophages. The infection assay was performed as in A. **E**) Confocal microscopy images of macrophages with EPEC bacterial cell on the membrane. Panels show images of the transmitted light (TL), stained nuclei (magenta), actin filaments (yellow), EPEC cells (cyan), and merged images. Scale bar: 10 µm. Error bars in B and C represent the mean ± SD of three biological replicates from a representative experiment out of three independent experiments. All data points from three independent experiments are shown in A and D. Statistical analysis was performed using Student’s t-test (ns: P > 0.05; *: P < 0.05; **: P < 0.01; ***: P < 0.001; ****: P < 0.0001).

In addition, we followed the activities of the EvgS/EvgA and PhoQ/PhoP systems by monitoring the promoter reporters of P*_safA_-gfp* and P*_mgrB_-gfp*, respectively, in the EPEC strain under the same growth condition (Figure 5B). The *mgrB* deletion resulted in a de-repression of PhoQ, thus an increase in the P*_mgrB_-gfp* reporter expression. We also observed a decrease in P*_safA_-gfp* expression due to a stronger inhibition by PhoP-P, which might not be applicable to other genes in the EvgA regulon (Figure 5B). Deletion of *safA* did not cause a significant change in P*_safA_-gfp* expression, but resulted in a marked decrease in PhoQ/PhoP reporter expression. This indicates that the absence of SafA does not substantially affect activation of EvgS by low pH, but disrupts signal transmission from the EvgS/EvgA system to PhoQ/PhoP, consistent with our observation in MG1655 strains. Under higher magnesium conditions (10 mM), similar but less pronounced differences among the three strains were observed in their interactions with macrophages (Figure 5A and S12). The expression of the GFP reporters (P*_safA_-gfp* and P*_mgrB_-gfp*) was lower under high magnesium conditions (Figure 5B), indicating the overall decreased activity of the EvgS/EvgA and PhoQ/PhoP signaling network.

To confirm the effect of *safA* and *mgrB* deletion on EPEC ingestion by macrophages, we performed complementation experiments under the low-magnesium condition. To maintain the activation dynamics, a plasmid carrying the *safA* gene under its native promoter was introduced to the Δ*safA* strain. The resulting over-complementation (due to plasmid copy number) led to an increased P*_mgrB_* activity (Figure 5C, left) and a significantly reduced engulfment by macrophages when compared to the wild type (Figure 5D, left). For the Δ*mgrB* strain, arabinose-induced expression of *mgrB* from a pBAD-driven plasmid restored both P*_mgrB_* activity (Figure 5C, right) and macrophage ingestion (Figure 5D, right) to the wild-type EPEC levels. Taken together, our findings indicate that the regulatory functions of both small proteins have physiological consequences, influencing EPEC evasion from phagocytosis with opposing effects. These phenotypes appeared to correlate with the level of PhoP regulon expression, and were more prominent under low-magnesium conditions.

EPEC employs distinct pathogenic strategies from *S. enterica*. Instead of replicating within macrophage phagosomes, EPEC is predominantly extracellular and may evade macrophage clearance by interfering with phagocytosis and modulating macrophage responses (57). To visualize EPEC–macrophage interactions, we labeled EPEC cells with mNeonGreen by introducing a plasmid expressing the fluorescent protein under a constitutive promoter. Following infection, macrophages were fixed and stained to visualize the actin filaments and the nuclei by confocal fluorescence microscopy. We observed macrophages with and without ingested EPEC (Figure S11C, z-stack movie S1 and S2) as expected. Interestingly, we captured another state of interaction, in which EPEC cells appeared to associate closely with the macrophage membrane, accompanied by local enrichment of actin filaments at the site of EPEC attachment (Figure 5E, z-stack movie S3). This observation raises the possibility that EPEC may influence macrophage engulfment through local modulation of the actin cytoskeleton. The physiological link between low pH-induced EvgS activation and EPEC evasion from macrophages remains unclear, and whether EvgS responds to other host-associated signals warrants further investigation. Nevertheless, our results reproducibly demonstrate that the activities of both two-component systems influence EPEC–macrophage interactions.

## Discussion

Our study of the small proteins (MgrB and SafA) revealed their distinct activation dynamics, magnesium-influenced interactions with PhoQ, and competitive binding. At low magnesium, *E. coli* cells respond to a low pH signal with an elevated activity of EvgA, a preferred binding of PhoQ to its activator SafA, and enhanced PhoQ activity, resulting in an augmented expression of both the PhoP and EvgA regulons. In comparison, high magnesium conditions dampen the bacterial response to low pH, demonstrated by a lower activity of EvgA, preferred binding of PhoQ to its inhibitor MgrB, and repressed PhoQ activity. Our findings establish magnesium as the central element to coordinate the opposing regulatory small proteins. This potentially enhances the sensitivity of *E. coli* to magnesium changes within a host, enables efficient responses to mild acidic pH, more intense stressors (such as strong acid and antimicrobials), and the host immune system via the signaling cascades from the EvgS/EvgA and PhoQ/PhoP systems.

A similar scenario of divalent cations that coordinate opposing regulatory small proteins has been reported for the cardiac SERCA pump in vertebrates (*48, 49*). SERCA transports Ca^2+^ into the sarcoplasmic reticulum and determines how quickly the heart muscle relaxes and refills with Ca²⁺ for the next contraction (*50*). In cardiac muscles, phospholamban (PLN) negatively regulates SERCA. The small protein DWORF opposes the PLN inhibition and activates SERCA (*48, 51, 52*). DWORF and PLN appeared to interact with SERCA at the same site (*52*) with Ca²⁺ coordinating their regulation. At high Ca²⁺ concentrations, calcium-bound SERCA changes conformation, which enhances the affinity of DWORF but reduces the binding of PLN, thereby promoting SERCA activation (*52*). At low Ca²⁺, the binding of PLN is favored, resulting in SERCA inhibition. This antagonistic regulation by PLN and DWORF enables the calcium-dependent adjustment of heart muscle contraction beat by beat. Mutation of PLN leads to the development of lethal dilated cardiomyopathy and heart failure in human (*50, 53*). Aside from their utilization of distinct divalent cations, the regulatory circuits of the SERCA/PLN/DWORF system exhibit a striking degree of similarity to the PhoQ/MgrB/SafA system in *E. coli*.

The *Salmonella* PhoQ is also regulated by additional proteins. Besides MgrB, the membrane protein UgtL interacts with PhoQ periplasmic and transmembrane domains, activating PhoQ autokinase activity at both neutral and acidic pH conditions (*54*). While UgtL is required for effective virulence, UgtS, another small protein, antagonizes the PhoQ activation by UgtL in *S. enterica*, and ensures the proper timing of virulence gene expression (*55*). The intricate regulation of PhoQ in *Salmonella* and *E. coli* underscores how PhoQ/PhoP are central regulators of virulence in these bacteria.

The additional layer of regulation by small proteins could improve bacterial adaptation to the host environment and limit energy expenditure on translation. In EPEC, deletion of *mgrB* resulted in a significantly reduced number of internalized bacteria. Although a slight increase in ingestion of the *safA* mutant was observed, this difference was marginal. Over-complementation of *safA* deletion resulted in a more marked difference in macrophage ingestion, confirming the effect of SafA on the interaction between EPEC and macrophages. Similar to our observation, Venturini, et al. (*56*) reported a reduced macrophage engulfment of an *mgrB* knockout in *S. enterica*, which they attributed to altered expression of motility and virulence genes. However, *Salmonella* and EPEC have divergent modes of pathogenesis. *S. enterica* is an intracellular pathogen that invades and replicates within macrophage phagosomes, whereas EPEC is primarily extracellular, typically degraded after uptake, and instead relies on inhibiting phagocytosis and altering macrophage responses (*57*). A possible explanation for EPEC evasion from macrophages involves PhoP regulation of the locus of enterocyte effacement (LEE) via the sRNA *mgrR* and the transcription regulator GrlA (*58–60*). LEE encodes the type III secretion system (T3SS) and associated effectors. Loss of MgrB enhances PhoQ activity, indirectly upregulating LEE genes, increasing T3SS expression, leading to inhibition of phagocytosis. In contrast, *safA* deletion disrupts activating signals to propagate from EvgS/EvgA to PhoQ/PhoP, likely resulting in increased macrophage engulfment of EPEC cells. While further experiments are required to verify this hypothesis, it is evident that the small proteins MgrB and SafA play a significant role in modulating EPEC virulence.

In this study, we revealed the existence of a regulatory factor that results in an elevated expression of the EvgA regulon in low magnesium conditions, suggesting an increased level of phosphorylated EvgA compared to that in high magnesium. EvgS, the cognate sensor kinase phosphorylating EvgA, responds to mildly acidic pH and redox cues (*41, 61*). Its activity can also be modulated by indole and nicotinamide through indirect or direct interactions (*62, 63*). To date, there is no evidence that EvgS directly senses magnesium. Consistent with this, EvgS lacks a membrane-proximal acidic surface, a structural feature characteristic of magnesium sensing in PhoQ (*64*). Therefore, it is unlikely that magnesium binds EvgS and affects its activity directly. However, it is possible that magnesium influences EvgS activity indirectly through other factors. The specific molecular mechanism of this magnesium-dependent regulation on EvgA activity remains to be determined.

While the regulatory network of SafA, MgrB and PhoQ are thoroughly studied here, the quantity of the native small proteins *in vivo* remains undetermined. Due to their short length and lack of trypsin digestion sites, they were not detectable with the proteomic approach in our attempts. The hydrophobicity makes them poor immunogens, consistent with the fact that validation of many small membrane proteins relies on epitope tagging followed by western blot (*5*). For MgrB and SafA, tagging at the cytoplasmic N-terminus does not significantly affect their function. However, this might influence their stability, since FtsH, a membrane-associated metalloprotease, recognizes cytosolic termini of integral membrane proteins, dislodges them, and then degrades these targets (*65*). Novel approaches are required to determine the native abundance of SafA and MgrB *in vivo*. Nevertheless, our findings revealed a sophisticated regulatory network, providing a scheme of a control module with self-tuning capabilities. Its small protein components represent promising candidates for novel therapeutic strategies.

## Material and methods

### Strains, plasmids and oligonucleotides

The *E. coli* strains, plasmids and oligonucleotides used in this study are listed in Tables S1 and S2.

### Bacterial growth conditions

*E. coli* cultures were cultivated in LB, minimal A medium (*66*), EvgS-optimized M9-minimal medium (*41*), or minimal E medium (*67*) as indicated. Media was supplemented with the appropriate antibiotics (100 µg/ml ampicillin, 34 µg/ml chloramphenicol and 50 µg/ml kanamycin) when applicable. Protein expression from pTrc99a was induced by 10 µM isopropyl β-D-1-thiogalactopyranoside (IPTG) and from pBAD33 by 0.008% arabinose, unless indicated otherwise. Liquid cultures were incubated at 37 °C with shaking at 200 rpm. To analyze bacterial growth over time, *E. coli* strains were grown overnight in the indicated growth medium, followed by 1:100 dilution into multi-well plates with the respective growth medium. Growth was monitored in an Infinite M Nano+ microplate reader (Tecan) at 37 °C by measuring absorbance at 600 nm every 15 min.

### GFP reporter assays

PhoQ activity was assessed using GFP reporter plasmids as described previously^35^. The required *E. coli* strains were transformed with a GFP reporter plasmid as indicated and additional plasmids as desired. Heterologous expression from pBAD33 and pTrc99a was induced as indicated in the figure legends. Cells were grown overnight in LB, washed with the respective minimal medium and then inoculated 1:100 in fresh medium. Stimuli were added to the medium at the start of the day culture. In experiments where cells were grown in minimal A medium, a second overnight culture was grown in minimal A medium. The cultures were grown Wuntil early log phase (OD = 0.4 to 0.5 for LB medium or OD = 0.15 – 0.2 for minimal medium) and fluorescence of cells was measured with a BD LSR Fortessa SORP flow cytometer (BD Biosciences) and the acquired data was analysed using BD FACSDiva software version 8.0 (BD Biosciences). Alternatively, cultures were grown in multi-well plates in a Tecan Infinite® 200 PRO plate reader and GFP fluorescence (excitation: 480 nm; emission: 510 nm) and OD_600_ were continuously measured for 24 h.

### Total RNA extraction and RT-qPCR

To quantify transcription of genes at different growth conditions, samples of culture were taken at desired time points and cell pellets were harvested. To monitor transcription dynamics over time after a switch from pH 7.5 to pH 5.5, *E. coli* MG1655 wild type was grown in EvgS-modified M9-minimal medium supplemented with 100 mM KCl at pH 7.5 to exponential phase. The cells were harvested and resuspended in EvgS-modified M9-minimal medium supplemented with 100 mM KCl at pH 5.5. To compare transcription rates between *E. coli* MG1655 wild type and *E. coli* MG1655 strains carrying genomic fusions of *phoQ*, *mgrB* and *safA* to mNeonGreen, cultures were grown in LB medium and EvgS-modified M9-minimal medium supplemented with 100 mM KCl at pH 5.5. The cultures were grown until early log phase (OD = 0.4 to 0.5 for LB medium or OD = 0.15 – 0.2 for minimal medium). Total RNA extraction and RT-qPCR was performed as described previously ^35^ with the difference that 2 ng/µl purified total RNA was used as template.

### Stimulus-dependent ratiometric FRET

*In vivo* interaction dynamics of PhoQ and MgrB or SafA under different conditions was analyzed by stimulus-dependent ratiometric FRET as described previously (*68*). An *E. coli* MG1655 Δ*phoPQ* Δ*mgrB* Δ*safA* strain carrying pBAD33_*phoQ*-mNeonGreen and either pTrc99a_mCherry-*mgrB* or pTrc99a_mCherry-*safA* was grown in LB medium supplemented with 10 mM MgSO_4_. Expression of the protein fusions was induced with 0.008 % arabinose (pBAD33) or 10 µM IPTG (pTrc99a). Cells were harvested and washed with tethering buffer (10 mM K_2_HPO_4_, 10 mM KH_2_PO_4_, 1 µM L-methionine, 10 mM soldium lactate, pH 7) supplemented with 10 mM MgSO_4_. Cells were attached to a glass coverslip using 0.1 % poly-L-Lysine. The coverslip was placed in a flow chamber that was mounted onto a dual-layer Nikon Ti-E inverted fluorescence microscope. The flow chamber was kept under a constant flow of 0.3 ml/min of tethering buffer. Stimuli were applied by a change of tethering buffer supplemented with the desired stimuli. The fluorescence signals of donor and acceptor were continuously recorded in the mNeonGreen and mCherry channels with an excitation time of 1 s. The donor was excited at 482/18 nm and its emission was detected at 525/50 nm, and the acceptor was excited at 585/29 nm while its emission was detected at 647/57 nm.

### Acceptor-photobleaching FRET

*E. coli* strains were grown and washed as described for stimulus-dependent ratiometric FRET. Additionally, plasmids pTrc99a_flag-flexi-mCherry-*mgrB*_flag-flexi-*safA* and pTrc99a_flag-flexi-mCherry-*safA*_flag-flexi-*mgrB* were used where indicated. Expression from pTrc99a plasmids was expressed using concentrations of IPTG as indicated.

For population-based acceptor-photobleaching FRET, cells were attached in a 96-well (Greiner) glass-bottom plate using 0.1 % poly-L-Lysine. Cells were overlaid with tethering buffer with the appropriate conditions to test. Acceptor photobleaching FRET was performed as described previously for the population-level with the exception of the use of a SOLA Lumencor light source (*36*). For single-cell measurements, the same filter configuration was used. Samples were illuminated using a 100×/1.45 NA Plan Apochromat objective. Images were acquired with a Hamamatsu Fusion sCMOS camera controlled by Nikon NIS-Elements software (version 5.XX). For each sample, four measurement points were selected. Image acquisition (1 s exposure time) was performed in the following sequence: (1) two images in the acceptor channel; (2) 110 images in the donor channel; (3) 7–14 s of acceptor photobleaching without image acquisition; (4) 50–70 images in the donor channel; and (5) two images in the acceptor channel. Image analysis was carried out as previously described (*36*).

For single-cell acceptor photobleaching FRET, cells were immobilized inside an Ibidi µ-slide 8 well glass bottom plate using agarose pads made from 1 % agarose and tethering buffer. For each sample, ten measuring points were selected and images were acquired with 1 s acquisition time as following: (1) 2 images in the acceptor channel; (2) 25 images in the donor channel; (3) 4 s of acceptor photobleaching (without image acquisition); (4) 25 images in the donor channel; (5) 2 images in the acceptor channel. Both for population- and single-cell level the acceptor photobleaching was performed in an area of 250 micron squared obtained by properly expanding the laser beam (ACAL BFI). The data analysis was performed using a custom pipeline comprising the following steps. (1) Channel registration and cropping were performed in Fiji to reduce image size for improved computational efficiency. (2) Cells were segmented on the fifth frame of the donor pre-bleaching sequence using CellPose (*69*) with the pre-trained model “bact_fluor_cp3” and a diameter parameter of 25. (3) Fluorescence intensities were quantified in Fiji for regions of interest with background subtraction in both bleached and unbleached areas. Donor intensity was calculated as the average of the last three frames before and the first three frames after acceptor photobleaching, while acceptor intensity was determined as the average of two frames acquired before and after photobleaching. The position of the bleached area was manually defined by the user and verified daily. (4) An output file containing all relevant parameters, including bleaching rate, FRET efficiency, donor and acceptor fluorescence intensities before and after bleaching, and cell areas were generated using a python script. FRET efficiencies were calculated taking into account the incomplete photobleaching of the acceptor, according to Equation 1 in the method published previously (*70*). Cells with a bleaching rate below 70% were excluded from analysis.

### Western-blot analysis of flag-tagged small proteins

Samples for western blot analysis of flag-tagged small proteins were denatured at 90 °C for 10 min. The samples were separated on a 16 % Tris-Tricine SDS gel (Thermofisher) using a SeeBlue™ Plus2 pre-stained protein standard (Invitrogen). Wet-tank transfer was performed using a PVDF membrane activated with methanol. Following transfer, the total protein was stained using a Revert™ 700 total protein stain kit. Monoclonal anti-FLAG M2 antibody was used as primary antibody and IRDye 800CW Goat anti-mouse IgG as the secondary antibody. The membrane was visualized using an Odyssey CLx imaging system (LI-COR) and bands were quantified with ImageJ.

### Overexpression and purification of recombinant mCherry and mNeonGreen

Recombinant mCherry and mNeonGreen fused to His-tags were overexpressed in *E. coli* BL21 (DE3). For this, the strains were grown in LB medium supplemented with the respective antibiotics until OD 0.4-0.6. Expression was induced with the addition of IPTG at a final concentration of 1 mM and growth was continued at 18 °C overnight shaking at 120 rpm. Cells were harvested and resuspended in lysis and binding buffer (100 mM NaH_2_PO_4_, 100 mM NaCl, 20 mM imidazole, 10 % glycerol, pH 7.8). The cells were disrupted using a sonicator at 80 % amplitude and a pulse of 0.5 s. Cell debris was removed by centrifugation at 3,220 rpm at 4 °C for 30 min and the cell lysate was filtered using 0.2 µm filters. The filtered cell lysate was loaded onto a HisTrap FF (1 ml) column pre-equilibrated with lysis and binding buffer on an Äkta pure (Cytiva) FPLC. After loading, the column was washed with lysis and binding buffer. The protein was eluted by applying a gradient from lysis and binding buffer to elution buffer (lysis and binding buffer with 300 mM imidazole). The protein peak was pooled and concentrated using Amicon Ultra-15 centrifuge filters (10 kDa). The concentrated protein was applied to a Superdex 200 Increase 10/300 GL (Cytiva) SEC column pre-equilibrated with the SEC buffer (20 mM NaH_2_PO_4_, pH 7.8, 50 mM NaCl). The protein was eluted by washing the column with the SEC buffer. The concentration of proteins was measured using a NanoDrop 2000 spectrophotometer (Thermo Scientific).

### Mathematical simulations of the PhoQ/PhoP and EvgS/EvgA signaling network

We model the entwined regulation of EvgS/EvgA, SafA, PhoQ/PhoP and MgrB expression as well as the activity of PhoQ as a dynamical system using a set of ordinary differential equation.

The PhoP-phosphorylation activity of PhoQ is modeled by assuming that PhoQ bound to MgrB is always inactive; PhoQ bound to SafA is always active, that MgrB and SafA bind competitively to PhoQ, and that in absence of small proteins, PhoQ is only inactive when bound to Mg^2+^. We also account for unequal dissociation constants of SafA and MgrB in low and high Mg^2+^ concentrations, assumed to reflect the Mg^2+^ unbound and bound states respectively, but we assume that the binding of Mg^2+^ to PhoQ is not affected by small proteins binding. All binding events follow first order kinetics. Under these assumptions, the fraction of phosphorylation-active PhoQ, which is also the fraction of phosphorylated PhoP is:

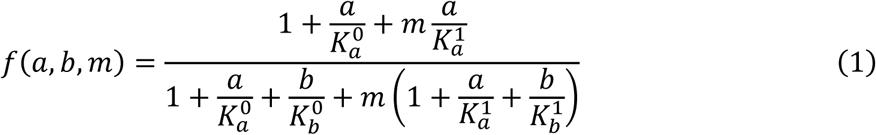

With *a* the SafA concentration, *b* the MgrB concentration, 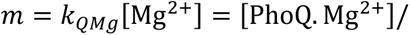 [PhoQ^*free*^] the normalized magnesium ion concentration, which reflects the level of magnesium-bound PhoQ, 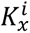 the dissociation constant for binding of species *x* to PhoQ in the magnesium bound (*i* = 1) or unbound (*i* = 0) state. We note that in the limit of large *a* and *b*, *f*(*a*, *b*, *m*) ≃ 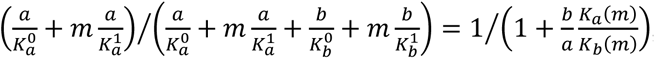, with 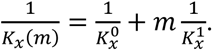. Hence, PhoQ activity only depends on *m* via the dissociation constants of SafA and MgrB in this limit.

The concentrations of the different species change due to induction and dilution by growth. Since PhoP and PhoQ are in a single operon, we assume that their concentrations are always proportional to each other, and, after normalization, equal to each other – this normalized concentration is noted *q*. We account for the induction of *safA* by EvgA-P, which we assume to be a constant throughout the simulation, as well as for the repression of *safA* and induction of *phoP/Q* and *mgrB* expression by PhoP-P, of normalized concentration *pp*. PhoP-P and EvgA-P are assumed to bind non-competitively to P*safA*. The negative feedback via YdeO on P*safA* is neglected (*41*).

The expression levels of SafA, PhoP/Q and MgrB are given by:

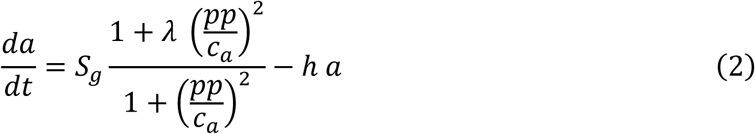

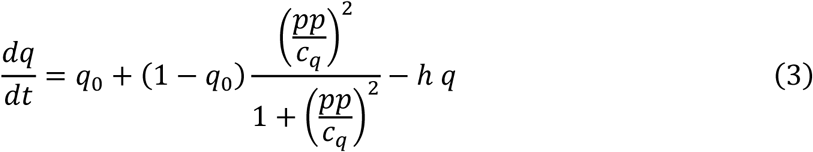

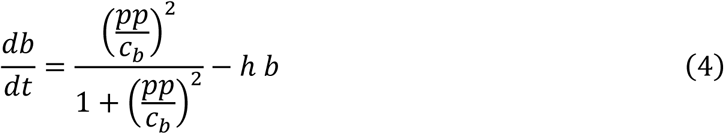

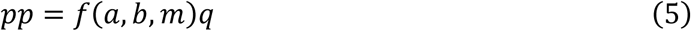

The dilution rate due to growth is ℎ = 0 in the lag phase and ℎ = 1 afterwards, so that time is normalized by the exponential growth rate of the cells. The concentrations of SafA and MgrB are normalized to the same factor, the maximal production of MgrB per unit of time, so that the normalized concentrations *a* and *b* are directly comparable. However, PhoP (PhoQ) concentration is normalized to its own maximal production per unit of time. The induction of *safA* upon EvgA-P promoter binding is accounted for by *S_g_*, which is the production rate of SafA upon induction by EvgA-P, normalized to the maximal production rate of MgrB. We used *S_g_* = 1 in the low magnesium concentration and *S_g_* = 0.65 when simulating reduced *safA* induction at high magnesium concentration. The pH=7 condition is modeled with *S_g_*= 0. The reduction factor of *safA* induction upon PhoP-P dimer binding is *λ* = 0.2. This value was chosen to reproduce the two-fold increase in *safA* induction upon PhoP deletion in the medium with [Mg^2+^] = 2 mM (*40*). We note *c_i_* the (normalized) binding constant of the PhoP-P dimer to the promoter of species *i*. The basal induction level of PhoQ/PhoP is *q*_0_ = 0.1. The quadratic dependences in *pp* reflect PhoP-P dimerization to bind promotors.

The dissociation constants for PhoQ binding were 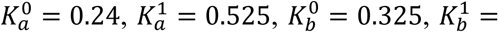 0.1, reflecting the measured dissociation constants (Fig 2D-G). The results were almost identical so long as the values of the dissociation constants were kept in the same ratios and were lower than the typical values for *a* and *b* during the simulation, i.e. PhoQ is saturated with SafA and MgrB. Much larger *K* values failed to capture the observed MgrB dynamics, indicating that SafA and MgrB competition for PhoQ binding likely occurs at saturation in experiments at pH=5. In the case where dissociation constants were taken independent of *m*, we used *K*=0.3 for the four affinities.

In [Mg^2+^] = 0.1 mM conditions, we calibrated *m_low_* = 0.4 using the data of Fig. 2D Δ*mgrB*, from which we deduce *m_high_* = 40 for [Mg2+] = 10 mM. For PhoP-P promotor binding, we used *c_q_* = 1.2 and *c_b_* = *c_q_*/4 = 0.3, to reflect the findings of Miyashiro and Goulian (*43*), accounting for the fact that our model has quadratic dependences in PhoP-P concentration, whereas they assumed a linear dependence in an effective PhoP-P concentration. Systematically varying it, we found dynamics that qualitatively matched experiments for *c_a_* > 0.15. We used *c_a_* = 0.4 in the simulations of Figure 4 and S9. Initial conditions were *a*(0) = 0, *b*(0) = 0.01, *q*(0) = *q*_0_. Any choice of small value for *b*(0) and *q*(0) gave very similar behavior.

We solve the set of equation by Euler integration using a time step *dt* = 0.01, which we checked to be sufficiently small for proper time integration. An initial lag phase of *T_lag_* = 3.4 (corresponding to about 5 doubling times t_1/2_) was followed by an exponential growth phase of the same duration, to mimic the experimentally observed behavior.

We additionally solved 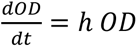 with *OD* (0) = 0.01 to generate the dependence of the protein concentrations on optical density.

### Whole-cell proteomic analysis

*E. coli* MG1655 cells were grown in minimal E medium at pH 5.5 supplemented with 0.1 mM or 10 mM MgSO_4_ at 37 °C with agitation. Cells were harvested at mid-log phase and washed three times with phosphate buffered saline (PBS). The resulting cell pellets were used for total protein extraction and proteomic analysis as described previously (*20*). The detailed procedure of proteomic analysis is described in the supplementary methods.

### Cultivation and differentiation of THP-1 monocytes

THP-1 human monocytic cells were cultured in RPMI-1640 medium supplemented with 10% heat-inactivated fetal bovine serum (FBS), 1 mM sodium pyruvate, and 1x GlutaMAX. For differentiation, cells were seeded at a density of 2.5×10⁵ cells per well in 24-well plates and treated with 20 nM phorbol 12-myristate 13-acetate (PMA). Plates were incubated for 24 h at 37 °C with 5% CO2 atmosphere for cells to differentiate. Successful differentiation was assessed by visually checking adherence to the bottom of the well and by observing the cell morphology.

### Infection of THP-1-derived macrophages

*E. coli* E611 wild type, Δ*mgrB* and Δ*safA* were grown in 5 ml LB medium overnight, washed and diluted 1:20,000 in minimal E medium at pH 5.5 with indicated magnesium concentrations. The strains were grown to mid-log phase in an Infinite® 200 PRO plate reader (Tecan). Bacteria were harvested by centrifugation (4,000 rpm, 10 min, room temperature) and resuspended in RPMI-1640 medium. A serial dilution (10⁻⁵ to 10⁻⁷) was plated on LB agar to determine the actual colony-forming units (CFU) input. Differentiated THP-1-derived macrophages were washed twice with PBS followed by adding fresh RPMI-1640 medium. The bacteria were added with a multiplicity of infection (MOI) of 100 in triplicates for each strain and time point, including negative controls (no bacteria). To facilitate contact between bacteria and macrophages, the plate was centrifuged at 700 × g for 10 minutes and incubated for 2 h at 37 °C and 5% CO₂.

The cells were washed twice with PBS and incubated for 1 h in RPMI containing 100 µg/ml gentamicin to eliminate extracellular bacteria. To quantify intracellular bacteria (time point 0 = 2 h post-infection), cells were lysed with 200 µl of 0.1% Triton X-100 in PBS. Released bacteria were serially diluted and plated on LB agar. The relative number of internalised bacteria was calculated as the mean percentage of CFUs recovered 2 h post-infection, normalized to the initial CFU input, which was defined as 100%.

### Confocal fluorescence microscopy

To visualize macrophages and EPEC cells during infection, THP-1-derived macrophages were seeded (1.25 × 10⁵ per well) in an 8-well μ-Slide (Ibidi GmbH). *E. coli* EPEC E611 strains were transformed with pLUC7 that expressing mNeonGreen under a constitutive promoter. The infection procedure was performed as described above without gentamicin treatment. Instead, the samples were fixed with 4% formaldehyde (v/v) for 20 min at room temperature, followed by three times washing with 1x PBS buffer. Triton X-100 (0.1% (v/v)) was then added and incubated for 30 min at room temperature to permeabilize cells. Samples were washed three times with PBS and blocked with 3% (w/v) bovine serum albumin (BSA) at 4 °C overnight to reduce background signal and non-specific staining. The samples were washed three times with PBS, incubated with ActinRed 555 (Invitrogen) with manufacturer recommended concentration for 20 minutes at room temperature in the dark, followed by one time PBS wash. The nuclei were stained with NucSpot 650/665 (Biotium, 1:1000 in PBS) for 15 minutes at room temperature in the dark, followed by one time PBS wash. Imaging was performed using a Zeiss LSM 880 laser scanning confocal microscope (Carl Zeiss Microscopy, Oberkochen, Germany). Images were acquired using a C-Apochromat 40×/1.2 W objective. Fluorescence signals were excited using laser lines at 488 nm, 543 nm and 633 nm. Z-stacks were acquired with a z-step size of 0.8 µm and 20–30 optical sections depending on the sample, to ensure complete coverage of the specimen. Imaging parameters were kept constant across all samples. ZEN (Blue Edition) software (Zeiss) was used for image processing.

## Supporting information

complete supplementary information

## Teaser

Two tiny bacterial proteins work together to fine-tune magnesium-dependent signaling that helps pathogenic *E. coli* evade immune cells.

## Acknowledgements

We thank Jörg Kahnt for performing mass spectrometry; Silvia González Sierra for technical assistance with flow cytometry; Lucas Cronau for the plasmid pLUC7; Dr. Lennart Randau (Philipps University Marburg) and Dr. Peter Graumann (Philipps University Marburg) for their suggestions concerning experiments; and members of the Yuan and the Sourjik laboratories for helpful discussions. AI-assisted editing tools were used to improve the readability of the text.

## Author contributions

JY, LW and VS designed the study; LW, GM, WL, JR, KHK and RC performed research; LW, WL, JR, KHK, GM, RC, TG and JY analyzed the data; and LW, WL, JR, GM, TG and RC wrote the first draft; JY, WL, GM, LW, KHK and RC edited and revised the manuscript.

## Data availability

The mass spectrometry proteomics data have been deposited to the ProteomeXchange Consortium via the PRIDE (*71*) partner repository with the dataset identifier PXD076996 (log in to the PRIDE website using the following details: project accession: PXD076996, token: ZnxCzcnzgcmh).

## References

1. S. Fuchs, S. Engelmann, Small proteins in bacteria – Big challenges in prediction and identification. Proteomics 23, (2023).

2. G. Storz, Y. I. Wolf, K. S. Ramamurthi, Small proteins can no longer be ignored. Annual review of biochemistry 83, 753–777 (2014).

3. A. T. Z. Burton, Rilee; Storz, Gisela, Large Roles of Small Proteins. Annual Review of Microbiology 78, (2024).

4. T. Gray, G. Storz, K. Papenfort, Small Proteins; Big Questions. Journal of Bacteriology 204, e00341–00321 (2022).

5. S. S. Yadavalli, J. Yuan, Bacterial Small Membrane Proteins: the Swiss Army Knife of Regulators at the Lipid Bilayer. Journal of bacteriology 204, e0034421 (2022).

6. R. Gelhausen et al., RiboReport - benchmarking tools for ribosome profiling-based identification of open reading frames in bacteria. Briefings in Bioinformatics 23, (2022).

7. C. H. Ahrens, J. T. Wade, M. M. Champion, J. D. Langer, A Practical Guide to Small Protein Discovery and Characterization Using Mass Spectrometry. Journal of Bacteriology 204, e0035321 (2022).

8. D. Remme, L. J. Tilg, Y. Pfander, J. Yuan, F. Narberhaus, Small DUF 1127 proteins regulate bacterial phosphate metabolism through protein-protein interactions with the sensor kinase PhoR. Microlife 6, uqaf023 (2025).

9. J. Yuan, H. G. Koch, B. A. Berghoff, Functional diversity and molecular interactions of small membrane proteins in bacteria. Microlife 6, uqaf035 (2025).

10. S. Cutting et al., SpoVM, a small protein essential to development in *Bacillus subtilis*, interacts with the ATP-dependent protease FtsH. Journal of Bacteriology 179, 5534–5542 (1997).

11. G. Karimova, M. Davi, D. Ladant, The β-Lactam Resistance Protein Blr, a Small Membrane Polypeptide, Is a Component of the *Escherichia coli* Cell Division Machinery. Journal of Bacteriology 194, 5576–5588 (2012).

12. E. Choi, K.-Y. Lee, D. Shin, The MgtR regulatory peptide negatively controls expression of the MgtA Mg2+ transporter in *Salmonella enterica* serovar Typhimurium. Biochemical and Biophysical Research Communications 417, 318–323 (2012).

13. A. M. Lippa, M. Goulian, Feedback inhibition in the PhoQ/PhoP signaling system by a membrane peptide. PLoS genetics 5, e1000788 (2009).

14. J. Yeom, Y. Shao, E. A. Groisman, Small proteins regulate *Salmonella* survival inside macrophages by controlling degradation of a magnesium transporter. Proceedings of the National Academy of Sciences of the United States of America 117, 20235–20243 (2020).

15. C. E. VanOrsdel et al., Identifying New Small Proteins in *Escherichia coli*. Proteomics 18, e1700064 (2018).

16. A. Kato, H. D. Chen, T. Latifi, Eduardo A. Groisman, Reciprocal Control between a Bacterium’s Regulatory System and the Modification Status of Its Lipopolysaccharide. Molecular Cell 47, 897–908 (2012).

17. E. A. Groisman, The pleiotropic two-component regulatory system PhoP-PhoQ. Journal of bacteriology 183, 1835–1842 (2001).

18. E. A. Groisman, A. Duprey, J. Choi, How the PhoP/PhoQ System Controls Virulence and Mg2+ Homeostasis: Lessons in Signal Transduction, Pathogenesis, Physiology, and Evolution. Microbiology and molecular biology reviews : MMBR 85, e0017620 (2021).

19. L. R. Prost et al., Activation of the bacterial sensor kinase PhoQ by acidic pH. Molecular cell 26, 165–174 (2007).

20. J. Yuan, F. Jin, T. Glatter, V. Sourjik, Osmosensing by the bacterial PhoQ/PhoP two-component system. Proceedings of the National Academy of Sciences of the United States of America 114, E10792–E10798 (2017).

21. E. G. Véscovi, F. C. Soncini, E. A. Groisman, Mg^2+^ as an Extracellular Signal: Environmental Regulation of *Salmonella* Virulence. Cell 84, 165–174 (1996).

22. M. W. Bader et al., Recognition of antimicrobial peptides by a bacterial sensor kinase. Cell 122, 461–472 (2005).

23. G. Viarengo et al., Unsaturated long chain free fatty acids are input signals of the *Salmonella enterica* PhoP/PhoQ regulatory system. The Journal of biological chemistry 288, 22346–22358 (2013).

24. I. Zwir et al., Dissecting the PhoP regulatory network of *Escherichia coli* and *Salmonella enterica*. Proceedings of the National Academy of Sciences of the United States of America 102, 2862–2867 (2005).

25. A. Kato, H. Tanabe, R. Utsumi, Molecular characterization of the PhoP-PhoQ two-component system in *Escherichia coli* K-12: identification of extracellular Mg2+- responsive promoters. Journal of bacteriology 181, 5516–5520 (1999).

26. Y. Eguchi, E. Ishii, M. Yamane, R. Utsumi, The connector SafA interacts with the multi-sensing domain of PhoQ in *Escherichia coli*. Molecular microbiology 85, 299–313 (2012).

27. E. A. Groisman, C. Parra-Lopez, M. Salcedo, C. J. Lipps, F. Heffron, Resistance to host antimicrobial peptides is necessary for *Salmonella* virulence. Proceedings of the National Academy of Sciences of the United States of America 89, 11939–11943 (1992).

28. Y. Liu et al., Magnesium sensing regulates intestinal colonization of enterohemorrhagic *Escherichia coli* O157: H7. MBio 11, 10.1128/mbio.02470-02420 (2020).

29. A. Platenkamp, J. L. Mellies, Environment Controls LEE Regulation in Enteropathogenic Escherichia coli. Frontiers in Microbiology 9, 1694 (2018).

30. H. W. Moon, S. C. Whipp, R. A. Argenzio, M. M. Levine, R. A. Giannella, Attaching and effacing activities of rabbit and human enteropathogenic *Escherichia coli* in pig and rabbit intestines. Infection and Immunity 41, 1340–1351 (1983).

31. A. S. Santos, B. B. Finlay, Bringing down the host: enteropathogenic and enterohaemorrhagic *Escherichia coli* effector-mediated subversion of host innate immune pathways. Cellular Microbiology 17, 318–332 (2015).

32. S. I. Miller, A. M. Kukral, J. J. Mekalanos, A two-component regulatory system (*phoP phoQ*) controls *Salmonella* typhimurium virulence. Proceedings of the National Academy of Sciences of the United States of America 86, 5054–5058 (1989).

33. T. Lemmin, C. S. Soto, G. Clinthorne, W. F. DeGrado, M. Dal Peraro, Assembly of the transmembrane domain of *E. coli* PhoQ histidine kinase: implications for signal transduction from molecular simulations. PLoS computational biology 9, e1002878 (2013).

34. M. E. Salazar, A. I. Podgornaia, M. T. Laub, The small membrane protein MgrB regulates PhoQ bifunctionality to control PhoP target gene expression dynamics. Molecular microbiology 102, 430–445 (2016).

35. S. Jiang et al., The inhibitory mechanism of a small protein reveals its role in antimicrobial peptide sensing. Proceedings of the National Academy of Sciences of the United States of America 120, (2023).

36. S. S. Yadavalli et al., Functional determinants of a small protein controlling a broadly conserved bacterial sensor kinase. Journal of bacteriology, (2020).

37. Y. Eguchi et al., B1500, a small membrane protein, connects the two-component systems EvgS/EvgA and PhoQ/PhoP in *Escherichia coli*. Proceedings of the National Academy of Sciences of the United States of America 104, 18712–18717 (2007).

38. E. Ishii, Y. Eguchi, R. Utsumi, Mechanism of activation of PhoQ/PhoP two-component signal transduction by SafA, an auxiliary protein of PhoQ histidine kinase in *Escherichia coli*. Bioscience, biotechnology, and biochemistry 77, 814–819 (2013).

39. Y. Eguchi, E. Ishii, K. Hata, R. Utsumi, Regulation of acid resistance by connectors of two-component signal transduction systems in *Escherichia coli*. Journal of bacteriology 193, 1222–1228 (2011).

40. N. A. Burton, M. D. Johnson, P. Antczak, A. Robinson, P. A. Lund, Novel aspects of the acid response network of *E. coli* K-12 are revealed by a study of transcriptional dynamics. Journal of molecular biology 401, 726–742 (2010).

41. Y. Eguchi, R. Utsumi, Alkali metals in addition to acidic pH activate the EvgS histidine kinase sensor in *Escherichia coli*. Journal of bacteriology 196, 3140–3149 (2014).

42. A. Zaslaver et al., A comprehensive library of fluorescent transcriptional reporters for *Escherichia coli*. Nature Methods 3, 623–628 (2006).

43. T. Miyashiro, M. Goulian, Stimulus-dependent differential regulation in the Escherichia coli PhoQ PhoP system. Proceedings of the National Academy of Sciences of the United States of America 104, 16305–16310 (2007).

44. H. Chen et al., Coordination of virulence factors and lifestyle transition in Pseudomonas aeruginosa through single-cell analysis. Communications Biology 8, 1236 (2025).

45. E. G. Vescovi, Y. M. Ayala, E. Di Cera, E. A. Groisman, Characterization of the bacterial sensor protein PhoQ. Evidence for distinct binding sites for Mg2+ and Ca2+. Journal of Biological Chemistry 272, 1440–1443 (1997).

46. L. Chen, L. Novicky, M. Merzlyakov, T. Hristov, K. Hristova, Measuring the energetics of membrane protein dimerization in mammalian membranes. Journal of the American Chemical Society 132, 3628–3635 (2010).

47. J. M. Autry et al., Oligomeric interactions of sarcolipin and the Ca-ATPase. Journal of Biological Chemistry 286, 31697–31706 (2011).

48. B. R. Nelson et al., A peptide encoded by a transcript annotated as long noncoding RNA enhances SERCA activity in muscle. Science 351, 271–275 (2016).

49. D. M. Anderson et al., Widespread control of calcium signaling by a family of SERCA-inhibiting micropeptides. Science Signal 9, ra119 (2016).

50. K. Haghighi et al., A mutation in the human phospholamban gene, deleting arginine 14, results in lethal, hereditary cardiomyopathy. Proceedings of the National Academy of Sciences of the United States of America 103, 1388–1393 (2006).

51. C. A. Makarewich et al., The DWORF micropeptide enhances contractility and prevents heart failure in a mouse model of dilated cardiomyopathy. Elife 7, (2018).

52. M. E. Fisher et al., Dwarf open reading frame (DWORF) is a direct activator of the sarcoplasmic reticulum calcium pump SERCA. Elife 10, (2021).

53. K. Haghighi et al., Human phospholamban null results in lethal dilated cardiomyopathy revealing a critical difference between mouse and human. Journal of Clinical Investigation 111, 869–876 (2003).

54. J. Choi, E. A. Groisman, Activation of master virulence regulator PhoP in acidic pH requires the *Salmonella*-specific protein UgtL. Science Signaling 10, eaan6284 (2017).

55. H. Salvail, J. Choi, E. A. Groisman, Differential synthesis of novel small protein times *Salmonella* virulence program. PLOS Genetics 18, e1010074 (2022).

56. E. Venturini et al., A global data-driven census of *Salmonella* small proteins and their potential functions in bacterial virulence. Microlife 1, uqaa002 (2020).

57. D. L. Goosney, J. Celli, B. Kenny, B. B. Finlay, Enteropathogenic Escherichia coli inhibits phagocytosis. Infection and Immunity 67, 490–495 (1999).

58. T. K. McDaniel, K. G. Jarvis, M. S. Donnenberg, J. B. Kaper, A genetic locus of enterocyte effacement conserved among diverse enterobacterial pathogens. Proceedings of the National Academy of Sciences of the United States of America 92, 1664–1668 (1995).

59. K. Moon, S. Gottesman, A PhoQ/P-regulated small RNA regulates sensitivity of Escherichia coli to antimicrobial peptides. Mol Microbiol 74, 1314–1330 (2009).

60. S. Bhatt et al., Hfq and three Hfq-dependent small regulatory RNAs-MgrR, RyhB and McaS-coregulate the locus of enterocyte effacement in enteropathogenic *Escherichia coli*. Pathogens and Disease 75, (2017).

61. S. Inada, T. Okajima, R. Utsumi, Y. Eguchi, Acid-sensing histidine kinase with a redox switch. Frontiers in Microbiology 12, 652546 (2021).

62. N. Boon et al., The signaling molecule indole inhibits induction of the AR2 acid resistance system in *Escherichia coli*. Frontiers in Microbiology 11, 474 (2020).

63. W. Yang et al., Enterohemorrhagic Escherichia coli senses microbiota-derived nicotinamide to increase its virulence and colonization in the large intestine. Cell Reports 42, (2023).

64. U. S. Cho et al., Metal bridges between the PhoQ sensor domain and the membrane regulate transmembrane signaling. Journal of Molecular Biology 356, 1193–1206 (2006).

65. K. Ito, Y. Akiyama, Cellular functions, mechanism of action, and regulation of FtsH protease. Annual Review of Microbiology 59, 211–231 (2005).

66. J. H. Miller, A short course in bacterial genetics : a laboratory manual and handbook for Escherichia coli and related bacteria. (Cold Spring Harbor Laboratory Press, Plainview, N.Y., 1992).

67. H. J. Vogel, D. M. Bonner, Acetylornithinase of Escherichia coli: partial purification and some properties. Journal of Biological Chemistry 218, 97–106 (1956).

68. S. Bi et al., Dynamic fluctuations in a bacterial metabolic network. Nature Communications 14, 2173 (2023).

69. C. Stringer, T. Wang, M. Michaelos, M. Pachitariu, Cellpose: a generalist algorithm for cellular segmentation. Nature Methods 18, 100–106 (2021).

70. J. Roszik, J. Szollosi, G. Vereb, AccPbFRET: an ImageJ plugin for semi-automatic, fully corrected analysis of acceptor photobleaching FRET images. BMC Bioinformatics 9, 346 (2008).

71. Y. Perez-Riverol et al., The PRIDE database at 20 years: 2025 update. Nucleic Acids Research 53, D543–D553 (2025).

