## Supplementary material for "Antagonistic small proteins enable magnesium-dependent tuning of two-component signaling": complete supplementary information

#### Supplementary methods

##### Construction of *E. coli* strains and plasmids

Genes in the genomic background of *E. coli* K-12 MG1655 (LW16, LW166), E611 (JR-S2-2, JR-S2-3) or BL21 (DE3) (LW156) were knocked out using lambda red recombination [1, 2]. For this, a kanamycin resistance cassette flanked by two FRT sites was amplified from pKD13 using primers with homologous arms for the gene of interest. Following insertion of the kanamycin resistance cassette into the genome using the lambda red recombinase expressed from pSIJ8, it was removed by FLP recombination using the FLP recombinase expressed from pSIJ8. The genomic fusions of *mgrB*, *safA* and *phoP* to *mneongreen* (LW287, LW289, LW291) were constructed in the background strain *E. coli* K-12 MG1655 using a similar approach as for the knockouts. However, the amplified kanamycin resistance cassette flanked by the two FRT sites was fused to the coding sequence for mNeonGreen and the primers introduced homologous regions for upstream or downstream of the gene of interest. The transcriptional reporter $P_{mgtLA}$-*gfp* was inserted into the coding sequence of *arsB* in the background strains *E. coli* K-12 MG1655 or *E. coli* K-12 MG1655 Δ*mgrB* (KH14, KH17) using a plasmid expressing NgAgo, an Argonaut protein, and encoding for the reporter and homologous sequences for the *arsB* locus (pKH2).

A plasmid encoding the transcriptional reporter $P_{safA}$-*gfp* (pLM01) was constructed by restriction digestion and ligation. The promoter region of *safA* was amplified from the genome of *E. coli* K-12 MG1655 using primers that included restriction sites for XhoI and BamHI. The resulting PCR product and the backbone pUA66 were digested using the respective restriction enzymes and ligated. Plasmids encoding for both *mgrB* and *safA* with fusions to *mcherry* and Flag-tags (pLW48, pLW49) were constructed using Gibson assembly by inserting a constitutive promoter and the coding sequence for the free small protein into pJY569 or pJY570. A plasmid encoding the first

60 bp of *malF* (corresponding to the first TM helix of the protein) fused to mCherry (pLW72) was constructed by using a pair of primers that each include one half of the desired sequence of *malF* and homologous sequences to amplify the pTrc99a backbone including mCherry from pJY569. Plasmids encoding for *mcherry* and *mneongreen* with a His-tag (pLW70, pLW71) were constructed using Gibson assembly. Sets of primers were used that amplified the backbone of pETDuet1 and the coding sequences of the fluorescent proteins from pJY569 or pJY573, respectively. A plasmid expressing NgAgo, an Argonaut protein, and encoding for the $P_{mgtLA}$-*gfp* reporter and homologous sequences for the *arsB* locus (pKH2) was constructed by Gibson assembly. Sets of primers were used to amplify the backbone of pB013 and the $P_{mgtLA}$-*gfp* reporter from the previous studies [3, 4].

**Quantification of intracellular fluorescent protein concentration**

To quantify the intracellular concentrations of mNeonGreen and mCherry in the strains used for FRET, the cells were harvested and resuspended in the SEC buffer. The cells were disrupted using a Precellys evolution homogenizer (Bertin). Cell debris was removed by centrifugation at 2,300 g for 10 min. mCherry and mNeonGreen fluorescence of lysed cells and purified fluorescent proteins was measured in an Infinite M200 Pro plate reader (Tecan) using excitation and emission wavelengths of 580 and 620 nm, and 497 and 527 nm, respectively. Intracellular mCherry and mNeonGreen concentrations were calculated by comparing the fluorescence of the lysed cells to the fluorescence of known concentrations of the purified fluorescent proteins.

**Proteomic analysis**

Proteins were isolated by resuspending cell pellets in 2% sodium lauroyl sarcosinate (SLS, in 100 mM ammonium bicarbonate) and heat exposure (90°C, 10 min). Protein concentration of debris-cleared lysates was measured by bicinchoninic acid protein assay (Thermo Scientific), followed by reduction with 5 mM Tris(2-carboxyethyl) phosphine (Thermo Fischer Scientific, 90°C for 15 min) and alkylation using 10 mM iodoacetamid (Sigma Aldrich, 20°C for 30 min in the dark). 50 µg protein was digested with 1 µg of trypsin (Serva) at 30°C over night. Post digest the SLS was precipitated by acidification and peptides were purified using C18 soli phase extraction.

Dried peptides were reconstituted in 0.1% trifluoroacetic acid and then analyzed using liquid-chromatography-mass spectrometry carried out on an Exploris 480 instrument connected to an Vanquish Neo and a nanospray flex ion source (all Thermo Scientific). The Vanquish Neo was operating in "Trap and Elute" mode using a Pepmap precolumn (Thermo Fisher) for peptide loading at 20 µl per min using solvent A (see below). The following separating gradient was used: 100% solvent A (0.1% formic acid) to 25% solvent B (99.85% acetonitrile, 0.15% formic acid) over 45 minutes, and an additional increase to 40% solvent B for 15 min at a flow rate of 300 nl/min.

MS raw data was acquired in data independent acquisition (DIA) mode. The funnel RF level was set to 40. Full MS resolution was set to 120.000 at m/z 200. AGC target value for fragment spectra was set at 3.000%. 67 windows of 6 Da were used between m/z 400-800. Resolution was set to 15.000 and IT to 22 ms. Stepped HCD collision energy of 22, 26, 30% was used. MS1 data was acquired in profile, MS2 DIA data in centroid mode.

Analysis of DIA data was performed using the DIA-NN [5] version 1.9 using a uniprot protein database from *E.coli* to generate a data set specific spectral library for the DIA analysis (full tryptic digest with two missed cleavage sites allowed, oxidized methionines (variable) and carbamidomethylated cysteins (fixed), N-term N extension). For DIA-NN search the following paramters were used: The precursor FDR = 1%; Mass accuracy, MS1 accuracy, Scan window set 0; Match between runs (MBR), Heuristic protein interference, No shared spectra set to 0; Protein interference = Off; Neural network classifier = Single-pass mode; Quantification strategy = QuantUMS (high precision); Cross-run normalization = RT-dependent; Library generation = IDs, RT & IM profiling; Speed and RAM usage = optimal results.

The DIA-NN report output was converted using an Rscript into aggregated protein information with peptide to protein assignment following the Occam´s razor approach. The output was further evaluated using the SafeQuant [6, 7].

#### Supplementary Figures

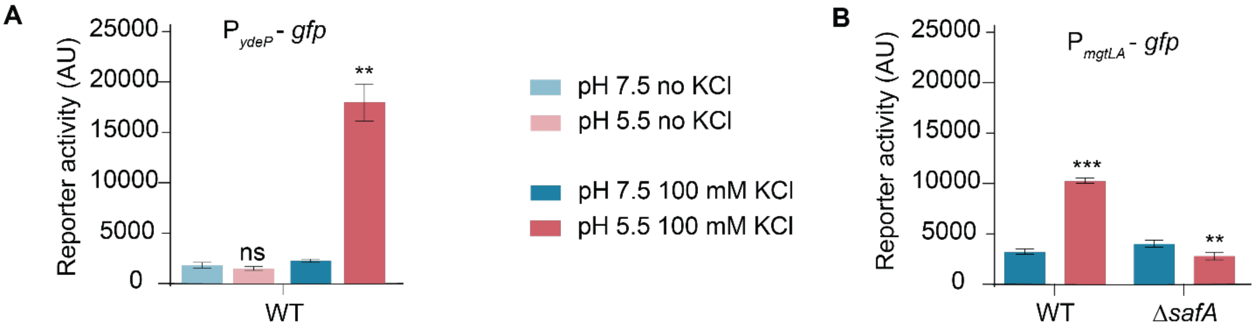


**Figure S1 Reporter assay of PhoQ and EvgS in M9 minimal medium.** Activity of PhoQ and EvgS in *E. coli* MG1655 in the presence (WT) and absence of SafA (Δ*safA*) using reporter plasmids carrying $P_{ydeP}$-*gfp* (**A**) or $P_{mgtLA}$-*gfp* (**B**). Strains were grown in M9 minimal medium at either pH 7.5 or 5.5 with or without the addition of KCl. Error bars represent the mean and standard deviation of the median fluorescence intensity of 30,000 cells of three biological replicates per condition. Shown is the representative result of one out of three independent experiments. P values were calculated using a Student's t-test (ns: P>0.05; *: P < 0.05; **: P < 0.01; ***: P < 0.001).

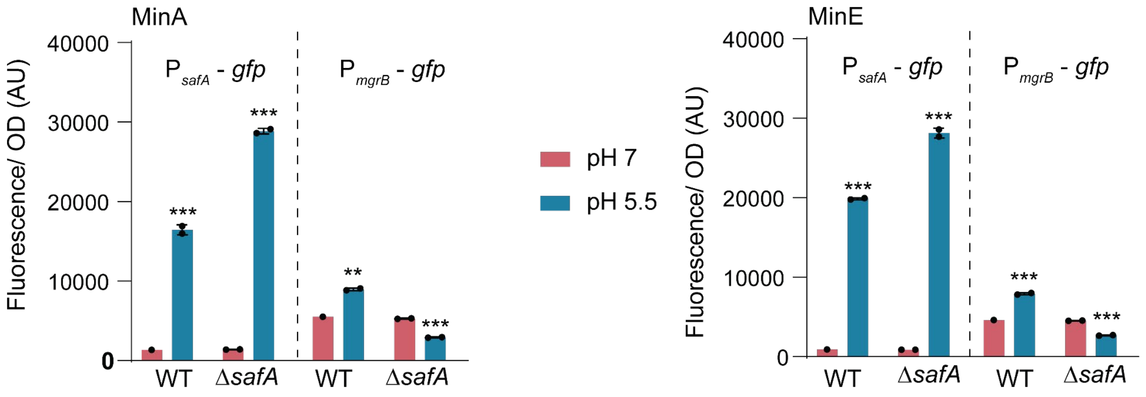


**Figure S2 GFP reporter activity in minimal A and E media.** *E. coli* MG1655 wild type and Δ*safA* strains carrying the indicated reporter plasmids were grown in minimal A (MinA) and minimal E (MinE) media at pH 7 or pH 5.5. The GFP fluorescence and $OD_{600}$ were measured in a plate reader at exponential growth phase. Error bars represent the mean and standard deviation of the median fluorescence intensity of 30,000 cells of two biological replicates per condition. Shown is the representative result of one out of two independent experiments. P values were calculated using a Student's t-test (ns: P>0.05; *: P < 0.05; **: P < 0.01; ***: P < 0.001).

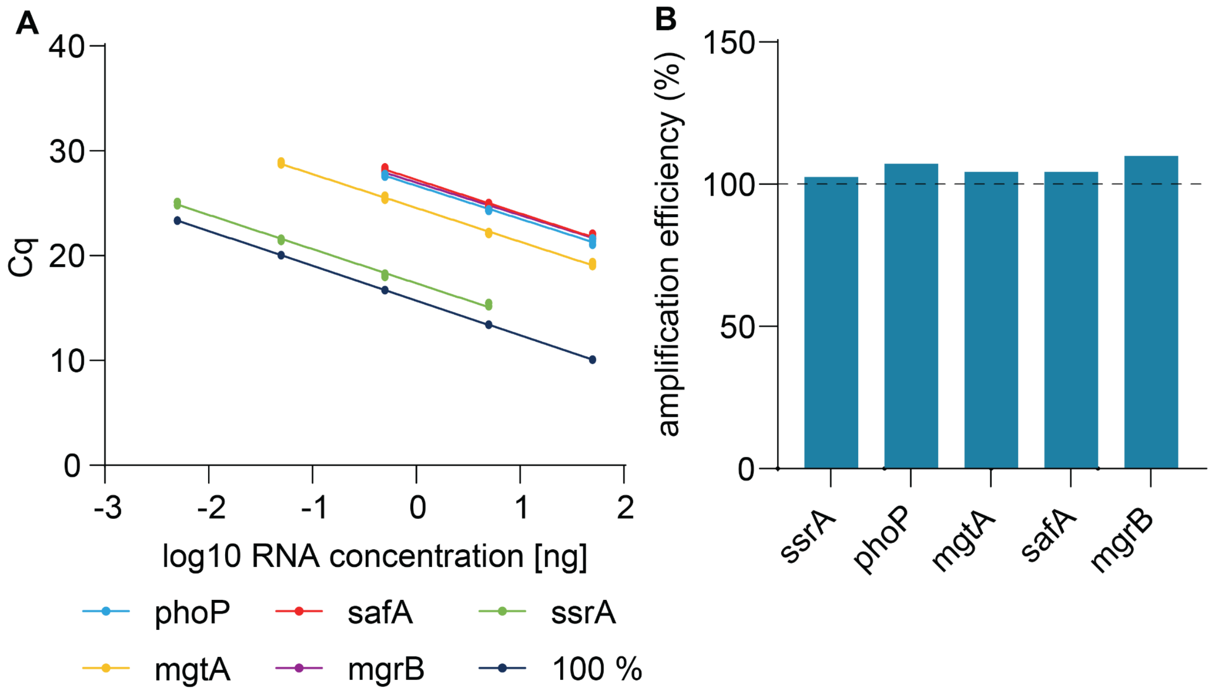


**Figure S3 Amplification efficiencies of primer pairs used for RT- qPCR.** Template RNA was prepared as serial dilutions of 10. Amplification efficiency was calculated from the slope of the decrease of Cq values over the log10 concentration of the template.

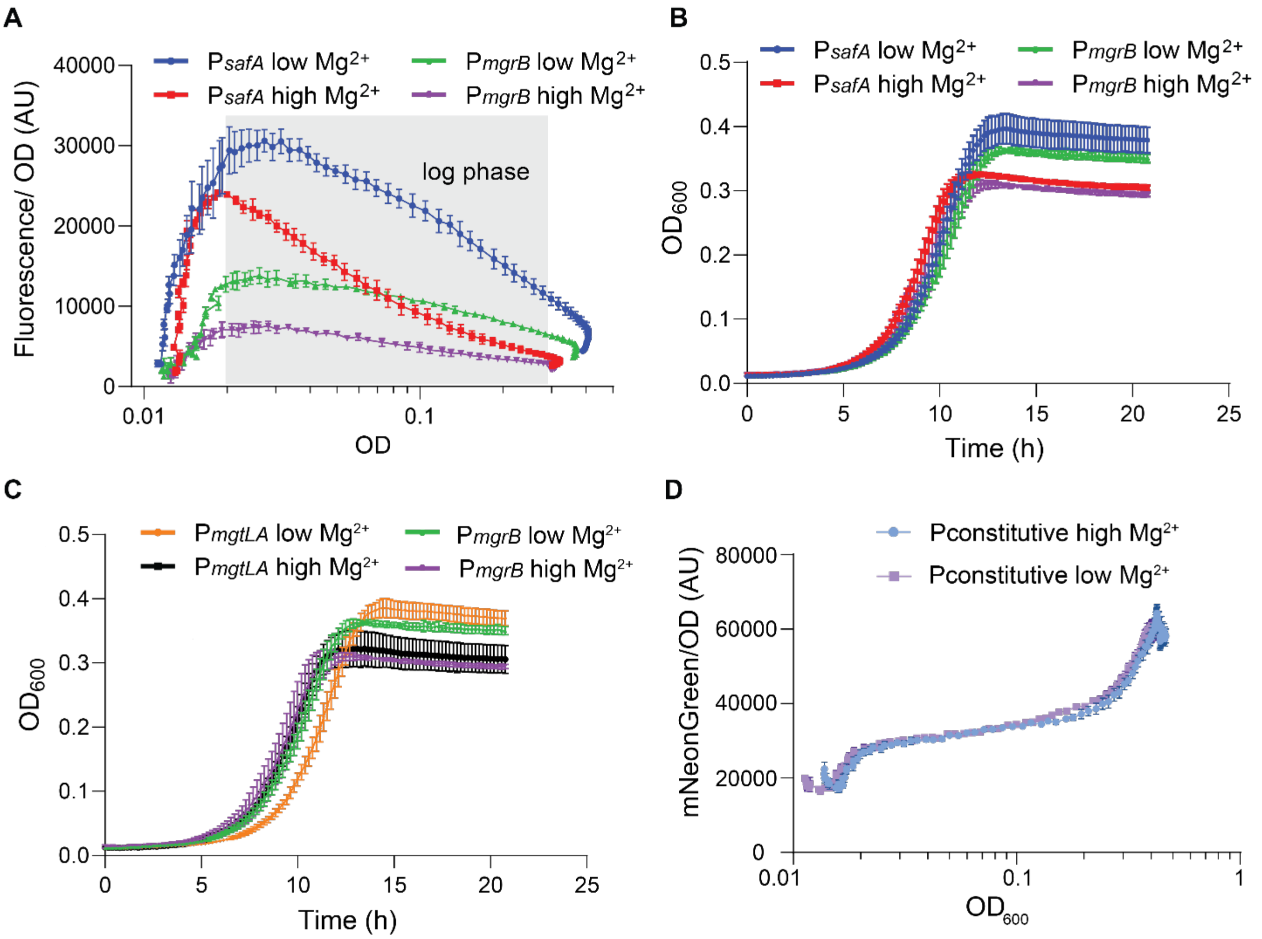


**Figure S4 Regulation of EvgS/EvgA and PhoQ/PhoP activities over time. A)** The activation profiles of *safA*, *mgrB* during the growth of the wild type cells in an EvgS activating medium with low or high magnesium concentrations. The wild type strain MG1655 was transformed with the respective GFP reporter plasmids ($P_{mgrB}$-*gfp*, and $P_{safA}$-*gfp*). GFP fluorescence and optical density (OD) were measured over the course of cell growth in minimal E medium with low (0.1 mM) or high (10 mM) $Mg^{2+}$ at pH 5.5. The ratio of GFP fluorescence to $OD_{600}$ were plotted over optical density. The grey-shaded area represents the exponential growth phase. **B, C**) The growth of MG1655 strain with indicated GFP reporters in minimal E medium with low (0.1 mM) or high (10 mM) $Mg^{2+}$ at pH 5.5. **D**) The expression of *mneongreen* under a constitutive promoter (pLUC7) was monitored during the growth of the wild type cells in an EvgS activating medium with low or high magnesium concentrations. Data points and error bars represent the mean and standard deviation of three biological replicates. One set of representative results out of three (in A, B, and C) or two (in D) independent experiments is shown.

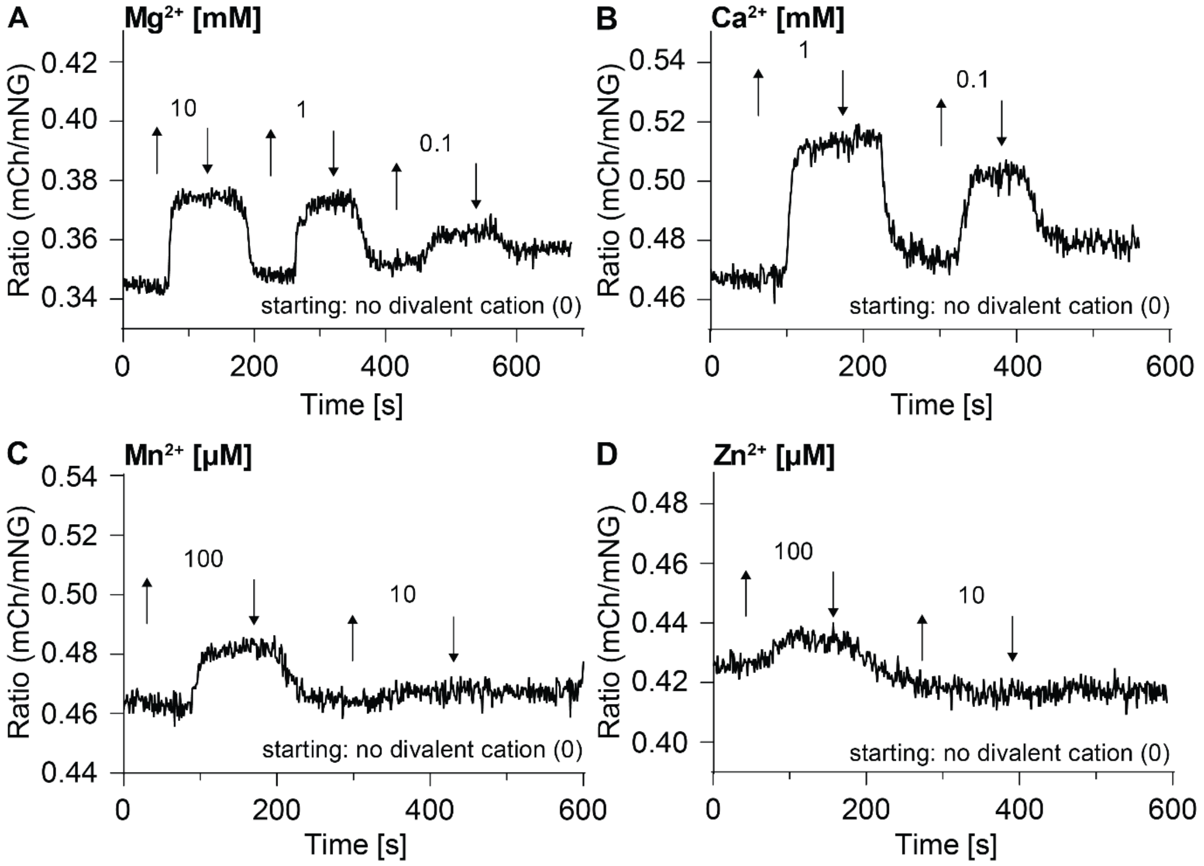


**Figure S5 The binding of MgrB to PhoQ in the presence of divalent cations.** *E. coli* MG1655 Δ*phoPQ* Δ*mgrB* Δ*safA* was transformed with pBAD33_*phoQ-mneongreen* and pTrc99a_*mcherry-mgrB*. Stimulus-dependent ratiometric FRET was performed by measuring mCherry and over mNeonGreen intensity with fluorescence microscopy. Divalent cations, $Mg^{2+}$ (**A**), $Ca^{2+}$ (**B**), $Mn^{2+}$ (**C**), and $Zn^{2+}$ (**D**) were added (upward arrow) and removed (downward arrow) at the indicated time points with the indicated concentrations in tethering buffers. Shown are the representative results of one out of three independent experiments.

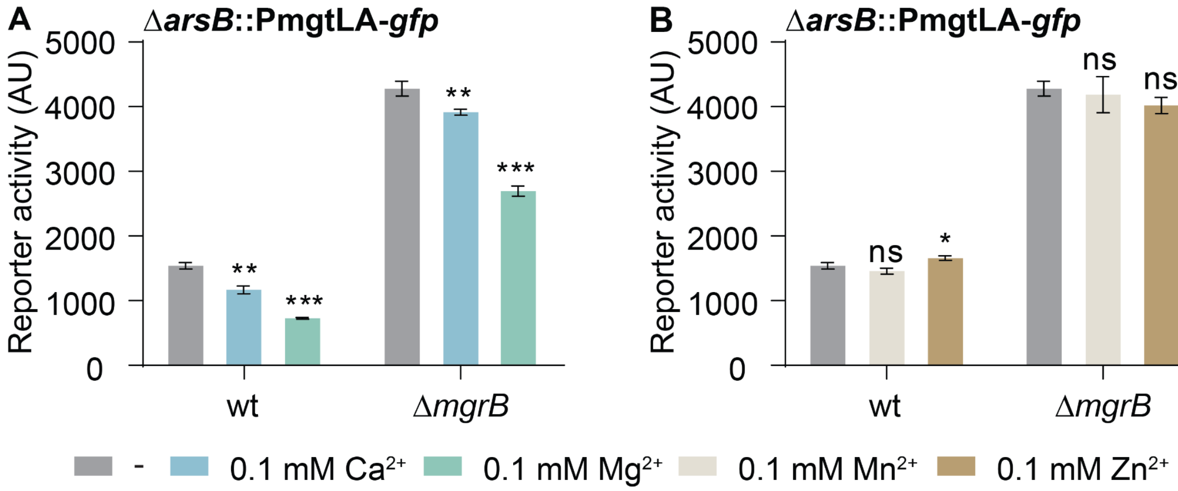


**Figure S6 PhoQ activity in the presence of divalent cations with or without MgrB.** PhoQ activity was monitored using the GFP reporter (Δ*arsB*::P$_{mgtLA}$-*gfp*) integrated into the genome of MG1655 wild type and *ΔmgrB* strains. Cells were grown in minimal A medium supplemented with the indicated concentrations of divalent cations. The GFP fluorescence were measured during the exponential growth phase. Error bars represent the mean and standard deviation of the median fluorescence intensity of 30,000 cells of three technical replicates per condition. Shown is the representative result of one out of three independent experiments. P values were calculated using a Student's t-test (ns: P>0.05; *: P < 0.05; **: P < 0.01; ***: P < 0.001).

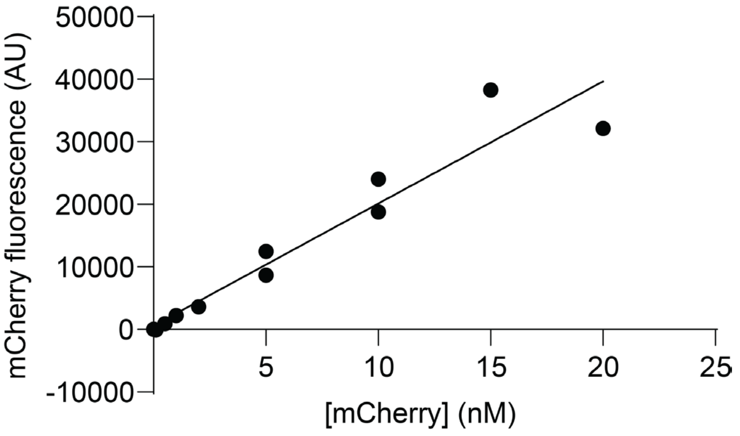


**Figure S7 Standard curve of mCherry fluorescence over mCherry concentration.** To convert intracellular mCherry fluorescence into copy number per cell, recombinant mCherry was purified and the fluorescence of different mCherry concentrations was measured in a plate reader to produce a standard curve.

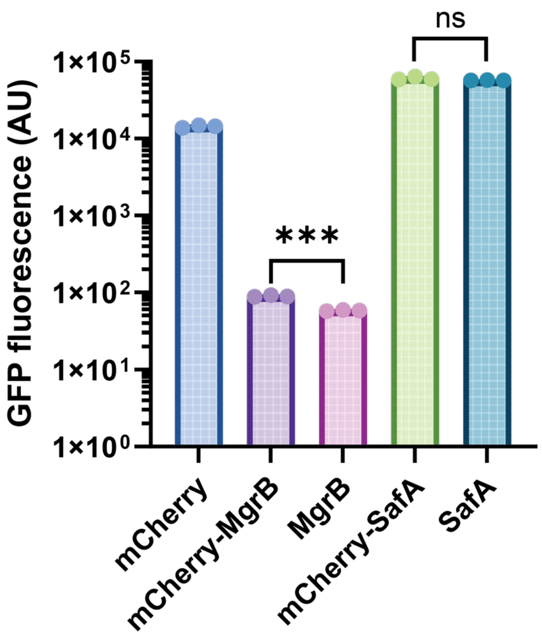


**Figure S8 The activity comparison of MgrB and SafA to their mCherry-fusions.** Activity of mCherry-fusions of MgrB and SafA was compared to non-fused versions by assessing PhoQ activity using the $P_{mgtLA}$ *-gfp* reporter. The proteins were expressed from plasmids in a Δ*mgrB* strain in LB. The strain with the expression of mCherry alone serves as a control. The mean and standard deviation of the median fluorescence intensity of 30,000 cells of three biological replicates is plotted. The representative result of two independent experiments is shown. P values were calculated using a Student's t-test (ns: P>0.05; *: P < 0.05; **: P < 0.01; ***: P < 0.001).

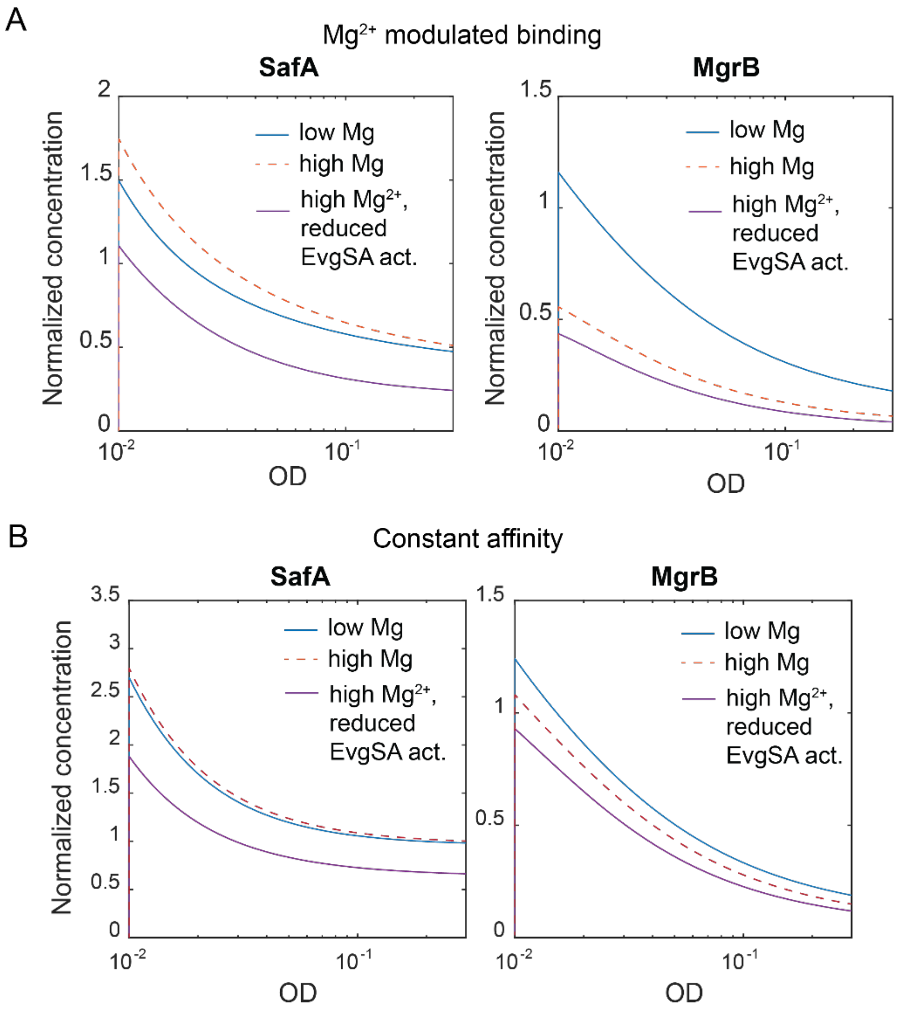


**Figure S9. Simulations of SafA and MgrB dynamics in high and low magnesium conditions upon stimulation with low pH.** The production of SafA was initiated by low pH stimulation of EvgS. The normalized production rate of SafA and MgrB was simulated over growth with (**A**) or without (**B**) the consideration of the magnesium-dependency regarding the affinity of the small proteins to PhoQ.

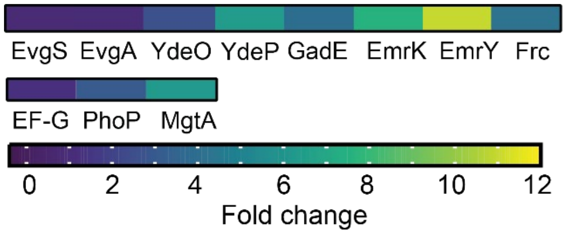


**Figure S10. Fold-change of protein abundance based on the proteomic data in Figure 4B.** The change in abundance of proteins regulated by EvgS/EvgA and PhoQ/PhoP is presented as a heat map.

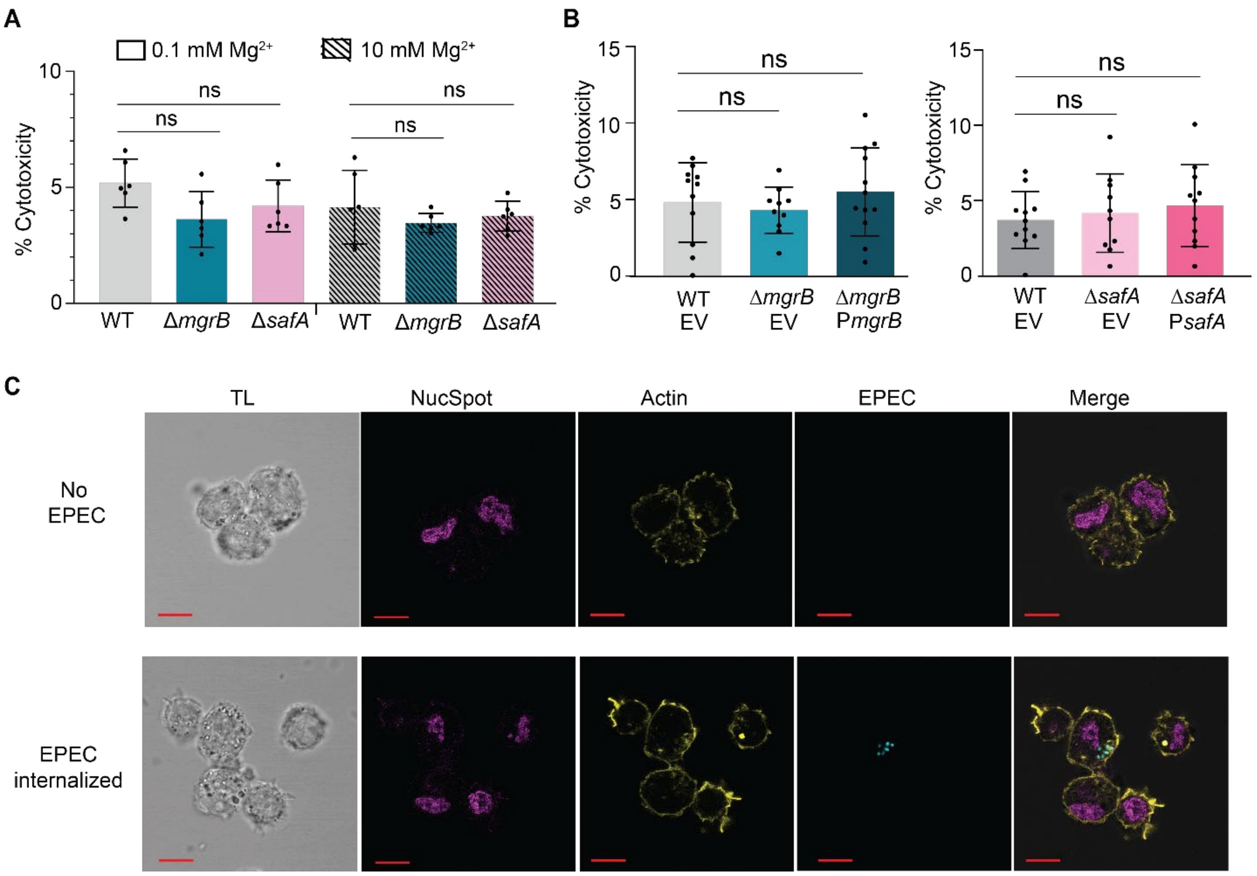


**Figure S11. THP-1-derived macrophages infected with EPEC E611 strains.** The cytotoxicity was measured by quantifying LDH release of THP-1-derived macrophages following infection with *E. coli* E611 wild type (WT), Δ*mgrB*, Δ*safA* (**A**) and the complemented strains (**B**) at time 1 h post infection. Data are shown as bar charts with calculated mean values and standard deviations. P values were calculated using a Student's t-test (ns: P>0.05; *: P < 0.05; **: P < 0.01; ***: P < 0.001). (**C**) The confocal microscopy images of macrophages with or without EPEC bacterial cell internalized. Panels show images of the transmitted light (TL), stained nuclei (magenta), actin filaments (yellow), EPEC cells (cyan), and merged images. Scale bar: 10 µm. The corresponding z-stack movies are deposited as supplementary files.

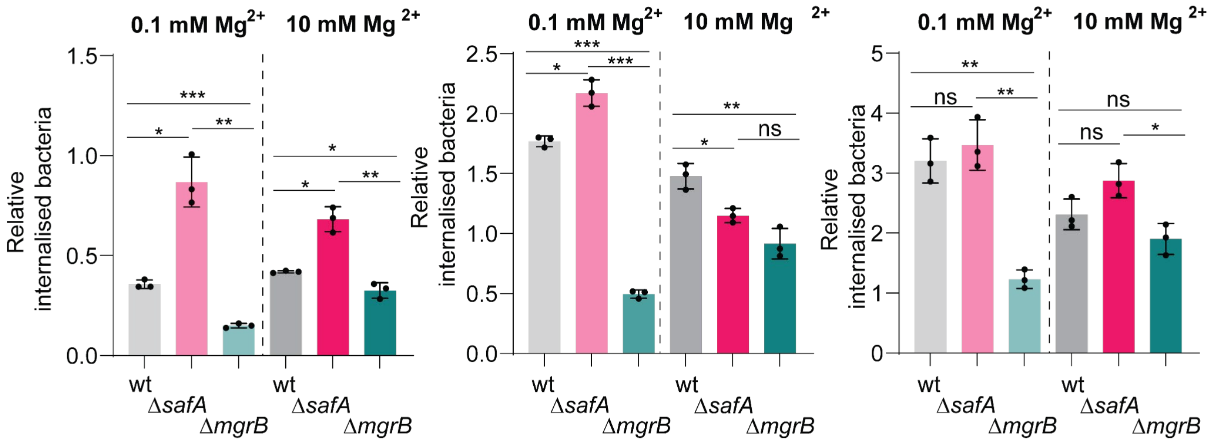


**Figure S12 Independent experiments of internalization of EPEC strains in macrophage infection assay**. Bacteria were grown in minimal E medium at pH 5.5 with either 0.1 or 10 mM $Mg^{2+}$ to log phase. Macrophages were infected with *E. coli* E611 at an MOI of 100 in RPMI medium for 2 h. Internalized bacteria were quantified. Each dot represents one of three biological replicates per strain. Statistical analysis was performed using a two-tailed unpaired Student's t-test assuming unequal variance (ns: $P > 0.05$; *: $P < 0.05$; **: $P < 0.01$; ***: $P < 0.001$).

#### Supplementary tables

#### Table S1. List of strains and plasmids used in this study

| Strain | Parental strain | Genotype | Source or reference |
|---|---|---|---|
| | DH5α | Wild type | Invitrogen |
| | MG1655 | Wild type | |
| LW16 | MG1655 | *ΔphoPQ ΔmgrB ΔsafA* | This study |
| LW166 | MG1655 | *ΔsafA* | This study |
| LW287 | MG1655 | *ΔmgrB::mneongreen-mgrB* | This study |
| LW289 | MG1655 | *ΔsafA::mneongreen-safA* | This study |
| LW291 | MG1655 | *ΔphoQ::phoQ-mneongreen* | This study |
| JingY15 | MG1655 | *ΔmgrB* | [4] |
| KH14 | MG1655 | *ΔmgrB ΔarsB::PmgtLA-gfp* | This study |
| KH17 | MG1655 | *ΔarsB::PmgtLA-gfp* | This study |
| LW156 | BL21 (DE3) | *ΔphoPQ* | This study |
| | E611 | Wild type | DSM no: 8701 |
| JR-S2-2 | E611 | *ΔsafA* | This study |
| JR-S2-3 | E611 | *ΔmgrB* | This study |

| Plasmid | Backbone | Insert | Source or reference |
|---|---|---|---|
| pSIJ8 | | flp_rhaS_rhaR | [2] |
| pKD13 | | FRT_KanR_FRT | [1] |
| pJingY361 | pUA66 | $P_{mgtLA}$-*gfp* | [4] |
| | pUA66 | $P_{ydeP}$-*gfp* | [8] |
| pLM01 | pUA66 | $P_{safA}$-*gfp* | This study |
| | pUA66 | $P_{mgrB}$-*gfp* | [8] |
| pJY573 | pBAD33 | *phoQ-mneongreen* | [9] |
| pOB2 | pTrc99a | *mcherry* | [9] |
| pJY569 | pTrc99a | *mcherry-mgrB* | [9] |
| pJY570 | pTrc99a | *mcherry-safA* | This study |
| pLW48 | pTrc99a | *trc-flag-flexi-mcherry-mgrB_sal-RBSimpr-flag-flexi-safA* | This study |
| pLW49 | pTrc99a | *trc-flag-flexi-mcherry-safA_sal-RBSimpr-flag-flexi-mrgB* | This study |
| pLW72 | pTrc99a | *mcherry-malfTM1* | This study |
| pLW70 | pETDuet1 | *mcherry-6His* | This study |
| pLW71 | pETDuet1 | *mneongreen-6His* | This study |

| | | | |
|---|---|---|---|
| pKH2 | | NgAgo_arsB_FRT_CamR_$P_{mgtLA}$-*gfp*_FR | This study |
| pKHK | p15A ori, CmR | $P_{safA}$-safA | This study |
| | pBAD33 | *mgrB* | [4] |

### Table S2. List of primers used in this study

| Primer | Sequence |
|---|---|
| mgrB-safA_Gibson_frag1_fw | GCAAGGCTGACGTTGCGAAGCAACGGCC |
| mgrB-safA_Gibson_frag1_rv | CATGAGAATTTTCAAATCCGCTCCCGGC |
| mgrB-safA_Gibson_frag2_fw | CGGATTTGAAAATTCTCATGTTTGACAGC |
| mgrB-safA_Gibson_frag3_rv | CTTCGCAACGTCAGCCTTGCAAACTATTG |
| Gibson_pLW29_fg2_rv_3 | CATCATCTTTATAATCCATATGGTACTCCTTATG |
| Gibson_pLW29_fg3_fw_3 | CATAAGGAGTACCATATGGATTATAAAGATGATG |
| Gibson_pLW32_fg1_fw | GTTTATCCCGTGGTGACGTTGCGAAGCAAC |
| Gibson_pLW32_fg3_rv | GTTGCTTCGCAACGTCACCACGGGATAAAC |
| Gibson_pLW32_fg0_fw | GTGGCGGAGGATCCATGGTGAGCAAGGG |
| Gibson_pLW32_fg0_rv | CCCTTGCTCACCATGGATCCTCCGCCAC |
| Q5mut_pLW72_fw | GTGGGTTACCTTGTTGTTTTAATGTACGCAAAGCTTGGCTGTTTTG |
| Q5mut_pLW72_rv | CAGCAGGCCGAGCAGACCTAGCACTGACCACATGAATACCTCCTTAAGTTG |
| Gibson_pLW70_i_fw | GAGATATACCATGGTGAGCAAGGGCGAG |
| Gibson_pLW70_i_rv | CGCCACCGCCCTTGTACAGCTCGTCCATGC |
| Gibson_pLW70_v_fw | GCTGTACAAGGGCGGTGGCGGTAGCCAT |
| Gibson_pLW70_v_rv | TGCTCACCATGGTATATCTCCTTCTTAAAGTTAAACAAAATTATTTCTAGAGGGGAATTGTTATCC |
| Gibson_pLW71_i_fw | GAGATATACCATGGTTTCTAAGGGTGAAG |
| Gibson_pLW71_i_rv | CGCCACCGCCCTTGTACAATTCGTCCATAC |
| Gibson_pLW71_v_fw | ATTGTACAAGGGCGGTGGCGGTAGCCAT |
| Gibson_pLW71_v_rv | TAGAAACCATGGTATATCTCCTTCTTAAAGTTAAACAAAATTATTTCTAGAGGGGAATTGTTATCC |
| Gibson_pKH2_i_fw | TGATTGCATGGTCTAAGAAACCATTATTATCATGAC |
| Gibson_pKH2_i_rv | GTCGACTGCGGCAAAACCCGTACCCTAG |
| Gibson_pKH2_v_fw | CGGGTTTTGCCGCAGTCGACGCGCAAAAAAC |
| Gibson_pKH2_i_fw | TTTCTTAGACCATGCAATCATGGTCTATATGAATATCCTCC |
| PsafA_XhoI_fw | TATCTCGAGTGCCTCTTCTCATTTCTTCTG |
| PsafA_BamHI_rv | TATGGATCCTTCAAATATGTTTATTTAGCGGATAAC |
| safA_H1P1 | ACTGATTAACGATTTTTAACGTTATCCGCTAAATAAACATATTTGAAATGATTCCGGGGATCCGTCGACC |
| safA_H2P2 | ATTTCATATTTATAATTTGCTGTTTGTTTTCAGCCTTGCAAACTATTGATTGTAGGCTGGAGCTGCTTCG |
| mgrB_H1P1 | AACGCATGCTAGTTTAATGACATAAGGTAGGTGAAACGGAGATTGGAGTGATTCCGGGGATCCGTCGACC |

| | |
|---|---|
| mgrB_H2P2 | GAAGAAAATCTAGTGCTGAAAAAATGATATCACCACGGGATAAACTGGTTTGTAGGCTGGAGCTGCTTCG |
| phoPQ_H1P1_BL21 | ATAATCGCGTTACACTATTTTAATAATTAAGACAGGGAGAAATAAAAATGATTCCGGGGATCCGTCGACC |
| phoPQ_H2P2_BL21 | TTAACGTAATGCGTGAAGTATGGACATATTTATTCATCTTTCGGCGTAGATGTAGGCTGGAGCTGCTTCG |
| mNG-mgrB-genomic_fw1 | GCTAGTTTAATGACATAAGGTAGGTGAAACGGAGATTGGAGTGAAAAAGTTTCGAGTTTCTAAGGGTGAAGAAGA |
| mNG-mgrB-genomic_fw2 | CCAGTTTATCCCGTGGTGATGATGATGAATTCCGGGGATCCGTCG |
| mNG-mgrB-genomic_rv1 | CGACGGATCCCCGGAATTCATCATCATCACCACGGGATAAACTGG |
| mNG-mgrB-genomic_rv2 | GAAGAAAATCTAGTGCTGAAAAAATGATATCACCACGGGATAAACTGGTTTGTAGGCTGGAGCTGCTTCG |
| mNG-safA-genomic_fw1 | CGATTTTTAACGTTATCCGCTAAATAAACATATTTGAAATGCATGCGACCACAGTTTCTAAGGGTGAAGAAGACAAC |
| mNG-safA-genomic_fw2 | CAATAGTTTGCAAGGCTGATGATGATGAATTCCGGGGATCCGTCGACC |
| mNG-safA-genomic_rv1 | GGTCGACGGATCCCCGGAATTCATCATCATCAGCCTTGCAAACTATTG |
| mNG-safA-genomic_rv2 | CATATTTATAATTTGCTGTTTGTTTTCAGCCTTGCAAACTATTGATTGTAGGCTGGAGCTGCTTCG |
| phoQ-mNG-genomic_fw1 | GGTGATTTTTGGTCGCCAGCATTCTGCGCCGAAAGATGAATCTAGAGTCGACGGCATGGTTTCTAAGGGTGAAG |
| phoQ-mNG-genomic_fw2 | GGACGAATTGTACAAGTAATGATGATGAATTCCGGGGATCCGTCGACC |
| phoQ-mNG-genomic_rv1 | GGTCGACGGATCCCCGGAATTCATCATCATTACTTGTACAATTCGTCC |
| phoQ-mNG-genomic_rv2 | GTAATGCGTGAAGTATGGGCATATTTATTCATCTTTCGGCGCAGATGTAGGCTGGAGCTGCTTCG |
| qPCR_safA_fw | CAAAATCACGCAAAGAGACAAC |
| qPCR_safA_rv | TGCGACCCATTTCTGAAATAC |
| qPCR_mgrB_fw | AGTTTCGATGGGTCGTTCTG |
| qPCR_mgrB_rv | TGATCGCACATCATGTTGAA |
| qPCR_mgtA_fw | GACCGTGATCGTGATGATTG |
| qPCR_mgtA_rv | CAGCCACGGGAAATAGCTTA |
| qPCR_phoP_fw | CCGGTGCTGATGATTATGTG |
| qPCR_phoP_rv | ATGACCTGTGAAGCCAGACC |
| qPCR_ssrA_fw | ATTCTGGATTCGACGGGATT |
| qPCR_ssrA_rv | AGTTTTCGTCGTTTGCGACT |
| Gibson_KHK67 | AGTTTGCAAGGCTGATCTAGAGTCGACCTGCAGGC |
| Gibson_KHK68 | GAGAAGAGGCATTAATTGGTAACGAATCAGACAATTGACG |
| Gibson_KHK69 | CTGATTCGTTACCAATTAATGCCTCTTCTCATTTCTTC |
| Gibson_KHK70 | CAGGTCGACTCTAGATCAGCCTTGCAAACTATTGATAATG |

#### References


1. Datsenko, K.A. and B.L. Wanner, *One-step inactivation of chromosomal genes in Escherichia coli K-12 using PCR products.* Proceedings of the National Academy of Sciences, 2000. **97**(12): p. 6640-6645.
2. Jensen, S.I., et al., *Seven gene deletions in seven days: Fast generation of Escherichia coli strains tolerant to acetate and osmotic stress.* Scientific reports, 2015. **5**: p. 17874.

3. Fu, L., et al., *The prokaryotic Argonaute proteins enhance homology sequence-directed recombination in bacteria.* Nucleic Acids Res, 2019. **47**(7): p. 3568-3579.
4. Yuan, J., et al., *Osmosensing by the bacterial PhoQ/PhoP two-component system.* Proceedings of the National Academy of Sciences of the United States of America, 2017. **114**(50): p. E10792-E10798.
5. Demichev, V., et al., *DIA-NN: neural networks and interference correction enable deep proteome coverage in high throughput.* Nat Methods, 2020. **17**(1): p. 41-44.
6. Glatter, T., et al., *Large-scale quantitative assessment of different in-solution protein digestion protocols reveals superior cleavage efficiency of tandem Lys-C/trypsin proteolysis over trypsin digestion.* J Proteome Res, 2012. **11**(11): p. 5145-56.
7. Ahrne, E., et al., *Critical assessment of proteome-wide label-free absolute abundance estimation strategies.* Proteomics, 2013. **13**(17): p. 2567-78.
8. Zaslaver, A., et al., *A comprehensive library of fluorescent transcriptional reporters for Escherichia coli.* Nat Methods, 2006. **3**(8): p. 623-8.
9. Yadavalli, S.S., et al., *Functional determinants of a small protein controlling a broadly conserved bacterial sensor kinase.* Journal of bacteriology, 2020.
